# Isoform-specific cofactor recruitment through the intrinsically disordered N-terminus of p63 underlies differential transcriptional activities

**DOI:** 10.64898/2026.04.19.719484

**Authors:** Marina F. Nogueira, Michael J. Moore, Aparna R. Biswas, Michael P. Meers, Sidharth V. Puram

## Abstract

The transcription factor p63 is critical for epithelial development and implicated in tumorigenesis. However, our understanding of the role of p63 in development and disease has been complicated by its diverse isoforms. As a member of the p53 family member of genes, *TP63* encodes for numerous isoforms, including the N-terminal variants TAp63 and ΔNp63, which are generated through alternative promoter usage. TAp63 and ΔNp63 share various structural domains, including the DNA-binding domain, and primarily differ in their N-terminus which consists of intrinsically disordered regions (IDRs). The isoforms are known to have different functions, including tumor suppression in the case of TAp63 and pro-tumor formation for ΔNp63, but how the N-terminus contributes to isoform-specific gene regulatory effects has yet to be elucidated. Using both genomic and TurboID proximity-labeling proteomic approaches, we show that the N-terminus mediates differential interactions with cofactors that have direct effects on isoform function, specifically the regulation of apoptosis. We find that the N-terminus of TAp63 interacts with more transcriptional machinery, leading to stronger transcriptional activity by TAp63 than ΔNp63. However, ΔNp63 maintains interactions with coactivators, suggesting it can retain some transactivation capabilities. Strikingly, the N-terminus of TAp63 displays enriched interactions with chromatin modifiers, including the histone acetyltransferase KAT2A, that result in TAp63-specific binding at inaccessible sites. We find that an IDR-mediated interaction with KAT2A is involved in regulation of apoptosis by TAp63. Collectively, our results suggest a model in which TAp63 and ΔNp63 broadly share genomic occupancy, but differential interactions with cofactors contribute to isoform-specific regulation by TAp63 and ΔNp63.

## INTRODUCTION

Transcription factors (TFs) regulate gene expression and play pivotal roles in development and disease. TFs are composed of a structured DNA-binding domain (DBD) that directs the TF to specific motifs in the genome, and effector domains, which mediate interactions with other TFs or cofactors. These effector domains can be folded into stable 3D structures or they can also consist of intrinsically disordered regions (IDRs). There has been significant progress in systematically characterizing and investigating the ≥1,600 TFs in the human genome (1), but due to technical limitations, many studies do not delineate how individual isoforms of a TF may influence gene expression. Given that 61% of TFs encode for multiple isoforms (2), this gap in isoform-specific knowledge represents a significant shortcoming in our understanding of human TFs.

p63 is a quintessential example of a TF in human development and pathophysiology for which isoform-specific differences remain poorly understood. A member of the p53 family, p63 is a critical TF in limb, craniofacial, and epithelial development (3, 4). However, our knowledge of this TF has been complicated by its numerous isoforms. Alternative splicing of the C-terminus can generate multiple isoforms, including ⍺, β, and *γ*;, while alternative promoter usage produces two classes of N-terminal variants with either a full-length transactivation domain (TAp63) or a truncated form (ΔNp63) (5).

In mouse models, the ΔNp63 isoforms have been shown to be the primary drivers of proper development (6), with TAp63 suggested to play a more modest, supportive role in epidermal formation (7). In cancer however, these two classes of isoforms appear to have opposing functions – TAp63 functions similarly to the classic tumor suppressor p53 (8–10), whereas ΔNp63 functions as an oncogene (11, 12). Accordingly, ΔNp63 is upregulated in squamous cell carcinoma (SCC), while expression of TAp63 is minimally detectable (13).

The C-terminal variants add an additional layer of complexity to regulation by p63 isoforms. TAp63⍺ and ΔNp63⍺ are believed to be the dominant isoforms expressed (13, 14), but other splice variants, like TAp63*γ*; and ΔNp63β, have increased transcriptional activity *in vitro* (5, 10, 15, 16). The full C-terminus is composed of an oligomerization domain (OD), sterile alpha motif domain (SAM), and an inhibitory domain (ID) (17). Alternative splicing can retain or remove these C-terminal domains in various combinations. Importantly, all isoforms of p63 share the same DBD, but how the different N-terminal and C-terminal effector domains contribute to divergent gene regulatory effects remains to be elucidated.

Here, we explore how the N-terminus of p63 encodes isoform-specific gene regulation and how these isoforms differentially regulate apoptosis. We found that both TAp63 and ΔNp63 transcriptionally activate target genes, but only TAp63 robustly induces apoptosis. TAp63 and ΔNp63 broadly exhibit similar binding across the genome, though we did identify isoform-specific peaks that were primarily attributed to TAp63 occupancy. However, differential binding did not explain differences in transcriptional regulation of apoptosis, prompting use of a proximity-labeling approach (TurboID) followed my mass spectrometry to identify differentially interacting cofactors involved in p63 regulation. By combining binding data, chromatin accessibility, and TurboID interactome data, we show that TAp63 neighbors more coactivators than ΔNp63 and its interactions with chromatin modifiers can explain TAp63-specific occupancy. We also demonstrate that an enriched, IDR-mediated interaction between TAp63 and the histone acetyltransferase KAT2A is critical for TAp63*γ*;-induced apoptosis. Collectively, our results elucidate how the N-terminus of p63 isoforms governs isoform-specific gene regulation, with implications for normal development and tumorigenesis. Our findings highlight the use of integrating multi-omic approaches to better understand individual TF isoforms and provide a roadmap for studying other important human TFs.

## MATERIALS AND METHODS

### Plasmid construction

The p63 isoforms TAp63α, TAp63γ, ΔNp63α, ΔNp63γ were cloned into the expression vectors described below. Importantly, we noticed a discrepancy between the literature and the TAp63α annotation in NCBI, which is annotated as transcript variant 1 (NM_003722.5), and TA* in the literature. For our experiments with TAp63, we utilized the CDS that begins at exon 2, which is consistent with what is used across the literature (5, 18, 19). For lentiviral, inducible expression of the p63 isoforms, we purchased the vector pLX401-INK4A (Addgene, plasmid #121919), removed the INK4A sequence, and cloned in the p63 isoform of interest. We either added a His-tag or a V5 tag to the C-terminus of the isoform. For the TurboID experiments, a pCDNA3.1 TurboID expression vector was kindly provided by Dr. Tim Ley (Washington University School of Medicine, St. Louis, MO). We further modified it to include a 15 amino acid linker sequence composed of glycine and serine between the TurboID and p63 sequences. TurboID was always fused to the N-terminus of the p63 isoform. For the control, we added a 3x nuclear localization signal (NLS) and the linker in front of the TurboID sequence. For tagged expression, each p63 isoform was cloned into a pCMV expression vector with a 3xFLAG sequence at the N-terminus. The Emerald GFP KAT2A plasmid (EmGFP-KAT2A) was purchased (Addgene, plasmid #65386). For all plasmid construction, Takara In-Fusion Snap Assembly was used according to the manufacturer’s protocols (Takara, 638945). All clones were verified by whole plasmid sequencing with Oxford Nanopore (Plasmidsaurus). Plasmids are available upon request.

### Cell lines

HNSCC cell lines PCI-30 (CVCL_C757) and SCC9 (CVCL_1685) were gifts from Dr. James Rocco (The Ohio State University, Columbus, OH). HEK293T were purchased from Takara (Takara Lenti-X, 632180). PCI-30 and 293T were cultured in high glucose, Dulbecco’s Modified Eagle Medium (DMEM) (Gibco, 11965118). SCC9 was cultured in Ham’s F-12 Nutrient Mixture (Gibco, 11765062) with DMEM at a ratio of 3:1. Cell culture media were supplemented with 10% fetal bovine serum (FBS) (Peak Serum, PS-FB1) and 1% penicillin-streptomycin (Pen-Strep) (Gibco, 15140122). For cell lines with integrated doxycycline-inducible vectors (Tetracycline-on vectors), media were supplemented with 10% Tet System Approved FBS (Takara, 631101) and 1% Pen-Strep. Cells were maintained at 37°C in a humidified 5% CO_2_ incubator. Cell lines were routinely tested for mycoplasma contamination.

### Lentivirus production and transduction of cell lines

LentiX HEK293T cells (Takara, 632180) were transfected with the appropriate plasmid and packaging vectors (psPAX2 and pMD2.G) at a ratio of 4:3:1 (ug of DNA). Viral supernatant was collected 48 hours later and sterile filtered. Cells were transduced with lentivirus supernatant in the presence of 8 ug/ml of polybrene (Santa Cruz Biotechnology 134220), followed by puromycin (2 ug/ml) antibiotic selection beginning 24 hours thereafter.

### *KAT2A* CRISPR-Cas9 knockout

For the *KAT2A* CRISPR knockout, we purchased lentiCRISPRv2 sgControl (Addgene, plasmid #125836) and lentiCRISPRv2 KAT2A sg2 (Addgene, plasmid #138186). The guide sequence was CACCgTCACCATGCCACCCTCAGAG. We transfected PCI-30 with the control plasmid and the *KAT2A* sg2 plasmid using Mirus TransIT-LT1 (Mirus, MIR2305). Following 24 hours, we added puromycin (2 ug/ml) to select for cells with the transfected vectors. Once all cells in the non-transfected control died, we stopped puromycin selection and expanded the cells. With Western blotting, we first verified that there was decreased KAT2A expression in the bulk population of cells. We then isolated single-cell clones by serial dilution. Editing was confirmed with targeted DNA sequencing (NGS) and Western blot.

### Pharmacologic agents

Doxycycline hyclate (Sigma-Aldrich, D9891) was reconstituted in water. GSK-699 PROTAC (InvivoChem, 2260944-68-9) and MG132 proteasome inhibitor (Selleck Chemicals, S2619) were reconstituted in DMSO (Sigma-Aldrich, D2650).

### Apoptosis Assays

PCI-30 and SCC9 cell lines transduced with specific pLX401-p63 isoforms were seeded at a density of 250–1,000 cells (depending on cell lines and days of assay) in 96-well plates in 200 ul medium. After a minimum of 24 hours, media was replaced and for the ‘+Dox’ conditions, media was supplemented with the titrated doxycycline (dox) concentration for that line (range 100 ng/ml to 0.5 ug/ml). Importantly, the ‘No Dox’ and ‘+Dox’ conditions for each line were always seeded on the same plate to control for technical variations. At the start of dox induction, one plate was used to determine base line levels of caspase-3/7 activation with the Caspase-Glo 3/7 Assay System (Promega, G8092). At the appropriate time intervals following dox induction (12 hours, 24 hours, etc.), we used the Caspase-Glo 3/7 Assay and measured luminescence with the BioTek Cytation 5 Cell Imaging Multimode Reader (Agilent). Relative caspase-3/7 activation was determined by the average luminescence across the technical replicates in the ‘+Dox’ condition divided by the average luminescence across the technical replicates in the ‘No Dox’ condition on the same plate. All functional assays were repeated with three to four independent experimental replicates.

### Western blotting

Cells were lysed on ice in RIPA buffer (Cell Signaling, 9806) plus a protease inhibitor cocktail (Sigma, P8340). The Pierce BCA Protein Assay Kit was used to determine protein concentration (Thermo Scientific, 23227). Cell lysates were mixed with 4x LDS buffer (Invitrogen, NP0007) and 100 mM DTT. Samples were run on Mini-PROTEAN TGX Precast SDS-PAGE gels (Biorad) in the Biorad Mini-PROTEAN Tetra Vertical Electrophoresis Cell. Gels were transferred onto PVDF membranes, blocked, and blotted using standard methods. Blots were incubated with primary antibodies overnight at 4°C followed by Horse radish peroxidase-linked secondary antibody (1:5000) incubation for 1 hour at room temperature. Blots were visualized using the iBright CL1000 Imaging System (Invitrogen).

### Antibodies

Primary antibodies used were: anti-p63 (Cell Signaling Technology CST, #39692), anti-V5 (CST, #13202), anti-His tag (CST, #2366), anti-DYKDDDDK (FLAG) tag (CST, #14793), anti-GFP (CST, #2956), anti-GCN5L2 (KAT2A) (CST, #3305), anti-PCAF (KAT2B) (CST, #3378), anti-BirA (Novus Bio, NBP2-59939), anti-β-actin (CST, #3700), anti-Puma (CST, #12450), anti-Noxa (CST, #14766), anti-cleaved PARP (CST, #5625), anti-Bcl2 (CST, #4223), anti-vinculin (CST, #13901). Secondary antibodies used were: anti-streptavidin HRP-linked (Invitrogen, S911), anti-rabbit IgG HRP-linked (CST, #7074), anti-mouse IgG HRP-linked (CST, #7076).

### RNA-sequencing

RNA was extracted using the RNeasy Mini Kit (Qiagen, 74104) with the on-column, DNA digestion protocol using RNase-free DNase I (Qiagen, 79254). The quality of extracted RNA was determined using RNA ScreenTape (Agilent, 5067-5576) and only RNA with RIN values >8.0 were used for sequencing. RNA was quantified by Qubit RNA XR (Invitrogen, Q33224) and 1 ug of RNA was used to generate libraries. Poly(A) selection was performed using NEBNext Poly(A) mRNA Magnetic Isolation Module (NEB E7490) followed by the Ultra II Directional RNA Library Prep Kit for Illumina (NEB, E7760). Sequencing was performed at the Genome Access Technology Center at the McDonnell Genome Institute (4444 Forest Park Ave., St. Louis, MO 63108) on an Illumina NovaSeq X Plus using paired end reads extending 150 bases. Basecalls and demultiplexing were performed with Illumina’s bcl2fastq software with a maximum of one mismatch in the indexing read. RNA-seq reads were then aligned to the Ensembl release 101 primary assembly with STAR version 2.7.9a1. Gene counts were derived from the number of uniquely aligned unambiguous reads by Subread:featureCount version 2.0.32. Isoform expression of known Ensembl transcripts were quantified with Salmon version 1.5.23. Sequencing performance was assessed for the total number of aligned reads, total number of uniquely aligned reads, and features detected. The ribosomal fraction, known junction saturation, and read distribution over known gene models were quantified with RSeQC version 4.04. Differential expression analysis was performed with the Python implementation of DESeq2 PyDESeq2 (20, 21).

### CUT&Tag and analysis

p63 CUT&Tag profiling was performed following the Benchtop CUT&Tag v3 protocol (22) to generate two replicates of each condition using the p63 and His-tag antibodies described in the primary antibodies section at a 1:25 dilution. Libraries were sequenced on an Illumina NovaSeq X Plus instrument (as described in the RNA-sequencing section) to obtain 150 bp paired-end reads. cutadapt (v 1.18) (23) was used to trim adapters. We aligned our reads with bowtie2 (v 2.3.5.1) (24) to hg38 using the parameters: --very-sensitive -local --no-unal --no-mixed --no-discordant --phred33 -I 10 -X 700. Unmapped, unpaired, and mitochondrial reads were removed using SAMtools (v 1.9) (25). PCR duplicates were removed using picard (v 3.0.0). We used deepTools (v 3.3.0) (26) bamCoverage with a bin size of 10 to create CPM normalized BigWig files and called peaks with MACS2 (q = 0.01) (v 2.1.2) (27). Peaks for biological replicates were combined using BEDTools (v 2.31.1) (28) intersect. To visualize peaks on the genome browser, replicates were then combined into a single BigWig file with deepTools bigwigCompare using the mean operation. To generate a matrix of peak counts, we used deepTools multiBamSummary. Heatmaps were generated using deepTools plotHeatmap. To annotate peaks with gene names and to find known motifs, we used HOMER (v 5.1) (29) findMotifsGenome.pl and annotatePeaks.pl.

### ATAC-sequencing and analysis

ATAC-sequencing was performed following the Omni-ATAC protocol (30). In brief, cells were harvested and pelleted by centrifugation at 1000g for 10 minutes at 4C. After removing the supernatant, cells were lysed using 100ul of ATAC lysis buffer (10mM Tris pH 7.4, 10mM NaCl, 3mM MgCl2, 0.1% NP40, 0.1% Tween-20, and 0.01% Digitonin) for 3min on ice after resuspension by pipetting. The lysis reaction was quenched with 1ml of resuspension buffer (10mM Tris pH 7.4, 10mM NaCl, 3mM MgCl2, 0.1% Tween-20) and inverted to mix. Nuclei were pelleted by centrifugation at 1000g for 10min at 4C and resuspended in 25ul 2x TD buffer (20mM Tris pH 7.6, 10mM MgCl2, 20% Dimethyl Formamide). Nuclei were counted using trypan blue and up to 50,000 nuclei per reaction were transferred to a new tube. Nuclei were filled to 12.5ul with 2x TD buffer and mixed with 12.5ul of transposition mix (1.25ul transposase, 8.25ul PBS, 0.25ul 1% Digitonin, 0.25ul 10% Tween-20, and 2.5ul H20). Transposition reactions were incubated on a shaking heat block at 37°C for 30min at 800rpm. DNA was purified using the Zymo DNA Clean and Concentrator kit (Zymo, D4004) with a final elution volume of 20ul. Final libraries were amplified using 9-11 PCR cycles and purified with Sera-Mag select beads (Cytiva, 29343052) using a double size selection with 0.55x and 1.5x sample volume beads (50ul PCR reaction volume). Libraries were sequenced on an Illumina NovaSeq X Plus instrument. Reads were aligned to hg38 with bowtie2 following adapter trimming with cutadapt. Unmapped, unpaired, and mitochondrial reads, along with ENCODE blacklist regions and PCR duplicates were removed as previously described in the CUT&Tag section. Peaks were called using MACS2 (q = 0.05). To visualize peaks, we used deepTools bamCoverage to generate RPGC normalized bigWig files. To call differential peaks, we used DiffBind (31) to generate a count matrix, and then used PyDESeq2 to find differentially accessible regions. We used HOMER findMotifsGenome.pl to find known motifs in the regions displaying increased accessibility as compared to the control.

### TurboID proximity labeling

PCI-30 cells were transfected with TurboID plasmids using Mirus TransIT-LT1 (Mirus Bio, 2305) transfection reagent. 24 hours following transfection, cell culture media was replaced with media containing 50 uM biotin (Milipore Sigma, 14400) and incubated for 12 hours. Following labeling, cells were collected and lysed in RIPA buffer (Cell Signaling, 9806) with P8340 protease inhibitor cocktail (Sigma-Aldrich, P8340). Lysates were sonicated using the Bioruptor Pico sonication device (Diagenode, B01080010) in 15ml Pico Tubes (Diagenode, C30010017) at 4°C for 5 cycles consisting of 30 seconds on, 30 seconds off, followed by brief vortexing of samples. Soluble lysate was collected and incubated with streptavidin resin (Pierce, 20361) at 4°C overnight on a rotator. The following day, beads were washed once with 1% SDS in PBS, three times with RIPA buffer, once with 50 mM Na_2_HPO_4_, 500 mM NaCl, 1% TritonX-100, pH 7.4, and once with PBS.

### Mass spectrometry

#### Peptide preparation

The peptides were prepared using a previously described method for on-bead tryptic digestion (32). The beads were washed four times with 1 ml of 50 mM ammonium bicarbonate buffer (pH = 8.0) (ABC) (Fluka, 09830). The washed beads were resuspended in 40 µl of ABC buffer containing 8 M urea (Sigma, U4884). The proteins were reduced by the addition of dithiothreitol (DTT) (Pierce, 20291), specifically 2 µl of 0.2 M DTT, and incubation for 60 min at 30 °C. The reduced proteins were alkylated using iodoacetamide (IAM) (Pierce, A39271) (1.7 µl of 0.5 M IAM) and incubation for 30 min at room temperature in the dark. The urea was diluted to 1.5 M by adding 160 µl of 50 mM ABC buffer prior to addition of 1 uAU lysyl endopeptidase (lys-C) (Wako Chemicals, 129-02541). Samples were digested for 2 hours at 30 °C in a Thermomixer with gyration at 750 rpm. 1 ug of sequencing grade modified trypsin (Promega, V5113) was added and the samples were incubated overnight at 30 °C in the Thermomixer gyrating at 750 rpm. The peptides were transferred to a new tube, the beads were washed with an additional 50 µl of ABC buffer and the wash was combined with the peptides. Residual detergent was removed by ethyl acetate extraction (33). Peptides were acidified to 1% (vol/vol) trifluoroacetic acid (TFA) (Sigma, 91707) in preparation for desalting using stage tips (C18) (34). The peptides were eluted with 60 µl of 60% (vol/vol) of acetonitrile (MeCN) (J.T. Baker, 9829-03) in 0.1% (vol/vol) formic acid (FA) (Sigma-Aldrich, 56302) and dried in a Speed-Vac (Thermo Scientific, Model No. Savant DNA 120 concentrator). The peptides were dissolved in 20 µl of 1% MeCN in water. An aliquot (10%) was removed for quantification using the Pierce™ Quantitative Fluorometric Peptide Assay kit. The remaining peptides were transferred to autosampler vials (Sun-Sri, 200046), dried and stored at −80°C for LC-MS analysis.

#### Ultra-high-performance liquid chromatography mass spectrometry – timsTOF

The peptides were analyzed using trapped ion mobility time-of-flight mass spectrometry (35). Peptides were separated using a nano-ELUTE chromatograph (Bruker Daltonics, Bremen, Germany) interfaced to a timsTOF Pro mass spectrometer (Bruker Daltonics) with a modified nano-electrospray source (CaptiveSpray, Bruker Daltonics). The mass spectrometer was operated in PASEF mode (35). Up to 200ng of sample in 2 µl of 1% (vol/vol) FA were injected onto a 75 µm i.d. × 25 cm Aurora Series column with a CSI emitter (Ionopticks). The column temperature was set to 50 °C. The column was equilibrated using constant pressure (800 bar) with 8 column volumes of solvent A (0.1% (vol/vol) FA). Sample loading was performed at constant pressure (800 bar) at a volume of 1 sample pick-up volume plus 2 µl. The peptides were eluted using one column separation mode with a flow rate of 300 nl/min and using solvents A (0.1% (vol/vol) FA) and B (0.1% (vol/vol) FA/MeCN): solvent A containing 2% B increased to 17% B over 60 min, to 25% B over 30 min, to 37% B over 10 min, to 80% B over 10 min and constant 80% B for 10 min. The MS1 and MS2 spectra were recorded from m/z 100 to 1700. The collision energy was ramped stepwise as a function of increasing ion mobility: 52 eV for 0–19% of the ramp time; 47 eV from 19–38%; 42 eV from 38–57%; 37 eV from 57–76%; and 32 eV for the remainder. The TIMS elution voltage was calibrated linearly using the Agilent ESI-L Tuning Mix (m/z 622, 922,1222).

#### Identification of Proteins

Unprocessed data from the mass spectrometer were converted to peak lists using Proteome Discoverer (version 2.1.0.81, Thermo-Fischer Scientific). The MS2 spectra from peptides with +2, +3 and +4 charge states were analyzed using Mascot software (Matrix Science, London, UK; version 2.8.1) (36). Mascot was set up to search the UNI-HUMAN-REF-plus_p63_isoforms_112823 database (unknown version, 20670 entries). The digestion enzyme was trypsin with a maximum of 4 missed cleavages allowed. The searches were performed with a fragment ion mass tolerance of 50 ppm and a parent ion tolerance of 25 ppm. Carbamidomethylation of cysteine was specified in Mascot as a fixed modification. Deamidation of asparagine, deamidation of glutamine, pyro-glutamate formation from n-terminal glutamine, acetylation of protein N-terminus and oxidation of methionine were specified as variable modifications. Peptides were filtered in Scaffold (version 5.2.1, Proteome Software Inc., Portland, OR) at 1% false-discovery rate (FDR) by searching against a reversed protein sequence database and a minimum of 2 peptides were required for protein identification.

### Spectral Count Analysis

Decoy and uncharacterized proteins were removed from the dataset. First, samples were normalized to the mean spectral count in the dataset. Specifically, counts were multiplied by the scaling factor i, where i = (mean total/total counts per sample). We then normalized all samples to the endogenously biotinylated Acetyl-CoA carboxylase (ACACA). Samples were normalized to the mean ACACA spectral count by multiplying the counts by scaling factor j, where j = (mean ACACA count in the dataset / ACACA count in specific sample).

For statistical analysis, proteins with normalized spectral counts < 2 in the isoform condition were omitted for further analysis. Log_2_ fold-changes (log_2_FC) between each isoform and the NLS-TurboID control were calculated using the mean spectral counts with a pseudocount of 0.001. Two-sample t-test with unequal variance were used to calculate p-values, followed by Benjamini-Hochberg FDR correction. High-confidence interacting proteins are defined as proteins with a log_2_FC > 0.6 compared to the NLS control and an adjusted p value (p_adj_) < 0.05.

### STRING Network Analysis

To look at the shared p63 interactome, we used the full STRING network (37) on shared proteins across isoforms that had a log_2_FC > 0.6 and p_adj_ < 0.05. The ‘textmining’, ‘neighborhood’, ‘experiments’, ‘databases’, ‘co-occurrence’, and ‘co-expression’ sources were all used to determine active interactions with a minimum required interaction score of 0.4. A functional enrichment of the network using STRING was also performed on the shared interactome. To look at the overall p63 interactome encompassing all four isoforms (i.e. 385 proteins, the union of interacting proteins), we looked at the physical network with a required interaction score of 0.9. Only ‘experiments’ and ‘databases’ were used for the interaction sources. We did not display unconnected nodes in the network.

To look at just the TAp63γ and ΔNp63γ interactomes separately, we looked at the physical network with a required interaction score of 0.7. Only ‘experiments’ and ‘databases’ were used for the interaction sources. We did not display unconnected nodes in the network.

### KAT2A co-immunoprecipitation

Briefly, 293T cells were transfected with EGFP-KAT2A plasmid and a pCMV-3xFlag vector containing the appropriate p63 isoform. Cells were collected 24 hours after transfection and lysed in 10 mM Tris-HCl (pH 8.0), 150 mM NaCl, 1mM EDTA, and 1% NP-40 along with P8340 protease inhibitor cocktail, 1 mM PMSF, 1 mM Na_3_VO_4_, and 10mM NaF. Lysates were incubated rotating with anti-DYKDDDDK (FLAG) magnetic beads (Pierce, A36797) at 4°C for 1-2 hours. Beads were subsequently washed five times with the same lysis buffer and eluted in Laemmli sample buffer under reducing conditions.

### IDR Prediction

Metapredict v3 was used to predict the per-residue disorder for each p63 isoform(38). We used the publicly available Google Collab notebook metapredict_batch.ipynb to perform batch predictions. The p63 DNA binding domain (DBD) was defined as the amino acid (aa) sequence encompassing the ten β-strands (S1 to S10) and three ⍺-helices (H1, H2, and H3) plus three amino acids before and after this region (39). The DBD corresponds to aa 135-325 for TAp63, and aa 80-270 for ΔNp63. We confirmed the DBD with AlphaFold (40, 41).

### Intermolecular Interactions Prediction

FINCHES was used to predict IDR-mediated interactions between p63 and KAT2A (42, 43). We utilized the FINCHES Google collab notebook to generate a .csv file of the interaction prediction values between p63 gamma (TA or ΔN) and KAT2A (Uniprot Q92830-1 canonical sequence was used for KAT2A) at each amino acid residue. We specifically used the Mpipi model with default parameters (mpipi_minmax 2.5, ticfreq 25, zero_folded True, window_size 31). Prediction values were plotted using matplotlib and seaborn.

## RESULTS

### TAp63 isoforms transcriptionally regulate apoptosis

To investigate the distinct effects of the p63 isoforms, we expressed TAp63⍺, TAp63*γ*;, ΔNp63⍺, and ΔNp63*γ*; in the head and neck SCC cell line PCI-30 (**Fig. 1A**). PCI-30 harbors a frameshift mutation in *TP53* and lacks endogenous expression of p63 and p73 (44, 45), making it an ideal system for isolating the effects of p63 isoforms, without confounding interactions from other p53 family members. The ⍺ isoforms are thought to be the predominant C-terminal variants expressed *in situ* (13, 14), but previous studies have shown that TAp63⍺ must be phosphorylated to be functional. Under conditions of stress, the inhibitory domain (ID) of TAp63⍺ is phosphorylated, which releases the AD from the ID, resulting in an open conformation that can form active tetramers (19, 46, 47). To avoid confounding effects of the phosphorylation and activation step, we focus on TAp63*γ*; since it lacks the ID and is active at baseline.

**Figure 1.**
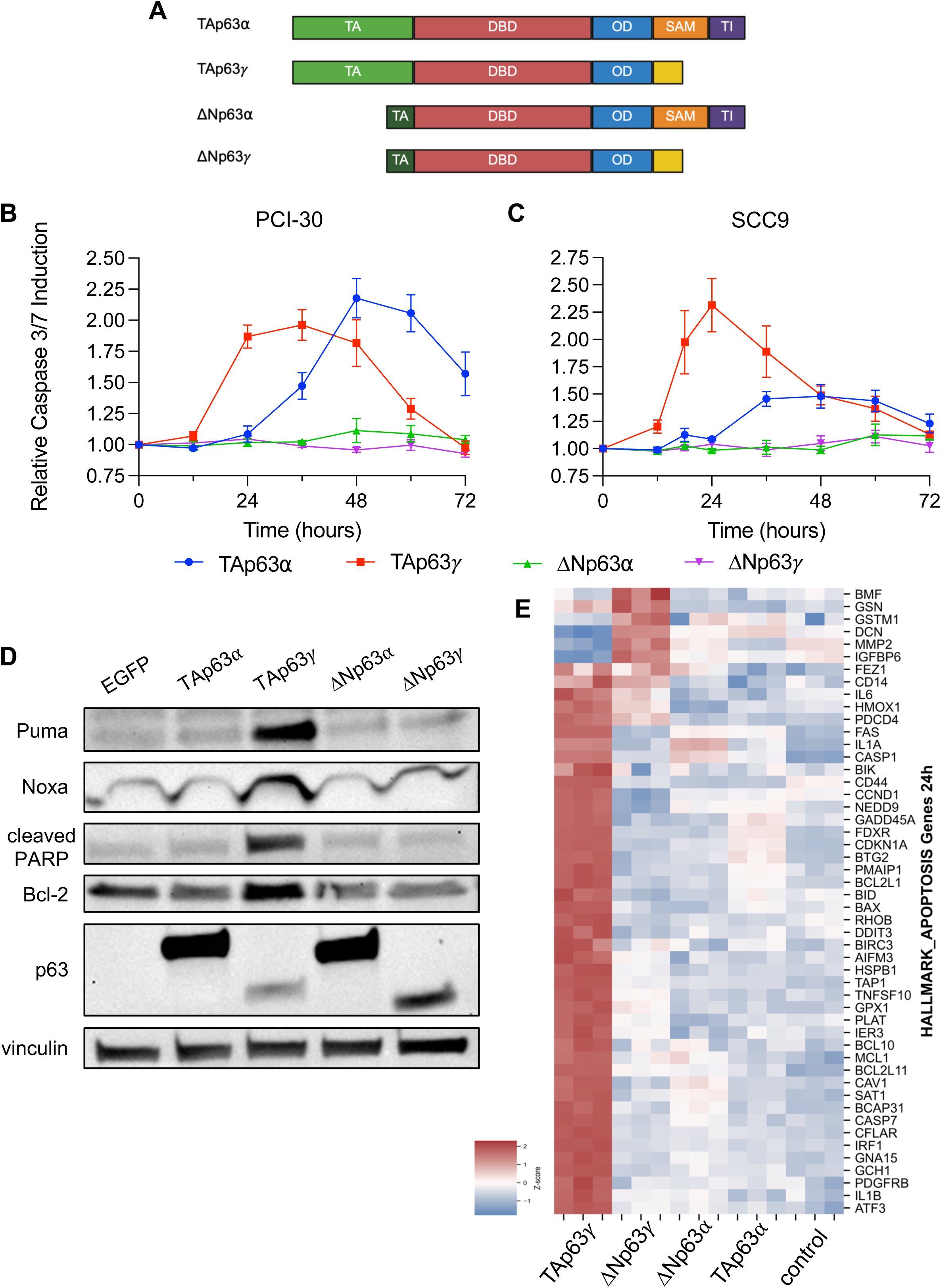
Regulatory effects of the p63 isoforms on apoptosis. **(A)** Schematic of p63 isoforms used in this study. Domains highlighted included the transactivation domain (TA), DNA-binding domain (DBD), oligomerization domain (OD), sterile alpha motif (SAM) domain, and the trans-inhibitory (TI) domain. **(B/C)** Relative luminescence measuring caspase 3/7 activation following expression of each p63 isoform in the cell line PCI-30 (B) or SCC9 (C). Measurements are relative to the no dox conditions for each cell line containing the dox-inducible lentiviral vector of the respective isoform. **(D)** Western blot of classic markers of apoptosis following expression of each p63 isoform. **(E)** Heatmap from RNA-sequencing of expression levels for genes in the MSigDB Hallmark apoptosis gene set following expression of each p63 isoform. Only genes that met a log_2_FC >1 and p_adj_ < 0.05 threshold for at least one isoform are displayed.

In order to look at the immediate and direct effects of p63 isoforms, we utilized a doxycycline-inducible lentiviral vector to individually overexpress each isoform. Doxycycline (dox) concentrations for induction were titrated for each vector to obtain comparable expression levels across all isoforms (**Fig. S1A**). However, we were unable to obtain similar expression of TAp63*γ*;. We observed that expression of TAp63*γ*; rapidly decreased over the span of 48 hours (**Fig. S1B**), suggesting the cells either promote degradation of TAp63*γ*; or undergo cell death. Treatment with the proteasome inhibitor MG-132 modestly increased TAp63*γ*; expression (**Fig. S1C**), indicating that reduced expression of TAp63*γ*; can be partially attributed to proteasomal degradation.

Upon expression of the p63 isoforms in cell culture, we immediately noted that cells expressing TAp63*γ*; do not survive. Similar to p53, TAp63 is known to regulate apoptosis (5). To quantify apoptosis, we measured caspase 3/7 activation through a luminescence assay following expression of each p63 isoform. In PCI-30, TAp63 isoforms activated caspases-3 and 7, while ΔNp63 had no effect (**Fig. 1B**). To validate these findings, we induced p63 isoform expression in the p63-null head and neck SCC cell line SCC9 (**Fig 1C**). In these cells, TAp63*γ*; also robustly induced caspase 3/7 activation, suggesting that this function of TAp63 is generalizable across cellular contexts.

To confirm TAp63 is initiating apoptosis through the intrinsic pathway, we examined protein expression of the BH3-only proteins PUMA and NOXA after inducing the p63 isoforms for 24 hours. We found that only TAp63*γ*; expression resulted in increased levels of pro-apoptotic PUMA and NOXA, as well as PARP cleavage (**Fig. 1D**). Previous studies have demonstrated that Bcl-2 blocks p53-dependent apoptosis (48, 49). To test the possibility that ΔNp63 isoforms are unable to promote apoptosis because they upregulate Bcl-2, we also examined Bcl-2 expression following p63 isoform induction. The ΔNp63 isoforms did not upregulate Bcl-2 (**Fig. 1D**), indicating that the difference in apoptosis induced by TAp63 versus ΔNp63 was not driven by changes in the balance of pro- and anti-apoptotic Bcl-2 family members.

Since the intrinsic apoptotic pathway is initiated by transcriptional regulation of pro-apoptotic factors, we sought to determine the transcriptomic effect of each p63 isoform. We performed RNA-sequencing following expression of each isoform for 24 hours. Notably, expression of the TAp63*γ*; isoform resulted in a greater number of differentially expressed genes (DEG) than the other isoforms (**Fig. S2A, S2B, S2C**). Consistent with TAp63*γ*; functioning as a constitutively active isoform of TAp63⍺, almost all genes regulated by TAp63⍺ were also regulated by TAp63*γ*; (**Fig. S2B, S2D**). Surprisingly, less than 20% of the DEG were shared by ΔNp63⍺ and ΔNp63*γ*; (**Fig. S2E**). These results indicate that unlike the TAp63 isoforms, the *γ*; C-terminal splice variant of ΔNp63 is not a more transcriptionally active variant than the ⍺ C-terminus. Rather, the C-terminus of the ΔNp63 isoforms contributes to unique transcriptional changes. Therefore, for our study, we chose to focus on TAp63*γ*; and ΔNp63*γ*; to avoid confounding C-terminal effects.

Apoptosis is a highly coordinated process that involves both transcriptional and signaling pathways, as well as a balance between pro- and anti-apoptotic factors. To confirm that TAp63 transcriptionally regulates apoptosis, we analyzed expression of genes found in the MSigDB HALLMARK apoptosis database, which consists of both positive and negative regulators of apoptosis. While the different p63 isoforms exhibit varying transcriptional effects on the MSigDB gene set, only TAp63*γ*; robustly increased the expression of these apoptosis-related genes (**Fig. 1E**). Together, our results indicate that while all isoforms of p63 can stimulate gene expression changes, only TAp63*γ*; is a strong transcriptional activator that robustly induces apoptosis.

### The N-terminus of TAp63 and ΔNp63 differ in acidity and length of predicted IDR

Given the observed differences between TAp63 and ΔNp63, we next asked how isoform-specific regulation might be encoded via differences in the N-terminal protein sequence. Interestingly, while N-terminal p63 variants have been extensively studied *in vivo*, it remains unknown how these variants function mechanistically at the transcriptional level. In particular, because the N-terminal p63 isoforms share an identical DBD, we were particularly interested in how such isoforms orchestrate disparate effects on the regulation of apoptosis. To explain the different transcriptional effects of TAp63 and ΔNp63 on apoptosis, ΔNp63 has typically been characterized as a repressor (5), but conflicting results in the literature and our own transcriptomic data indicate that ΔNp63 can also increase gene expression. Thus, despite a truncated N-terminus, there is evidence that ΔNp63 retains a small activation domain (50), raising the possibility that increases in gene expression could be a direct transcriptional effect.

ADs of TFs are known to be disordered. Recently, there has been significant work demonstrating the importance of intrinsically disordered regions (IDR) in TF gene regulation and TF binding preferences (56–58). To determine if the N-terminus of TAp63 and ΔNp63 are disordered, we used metapredict v3 to analyze the amino acid sequences of the isoforms (38). We found that despite the N-terminal truncation of ΔNp63, both ΔNp63 and TAp63 are predicted to be disordered (**Fig. 2A**). There is a clear decrease in the predicted disorder score at the boundaries around the DBD, with additional disordered regions predicted within the C-terminus. We also observed a decrease in predicted disorder in the first 10-20 amino acids of TAp63 that correspond to an alpha-helix. 51% of TAp63*γ*; is predicted to be disordered, while 44% of ΔNp63*γ*; is predicted to be disordered. Notably, ΔNp63 only contains 15 unique amino acids in the N-terminus compared to TAp63.

**Figure 2.**
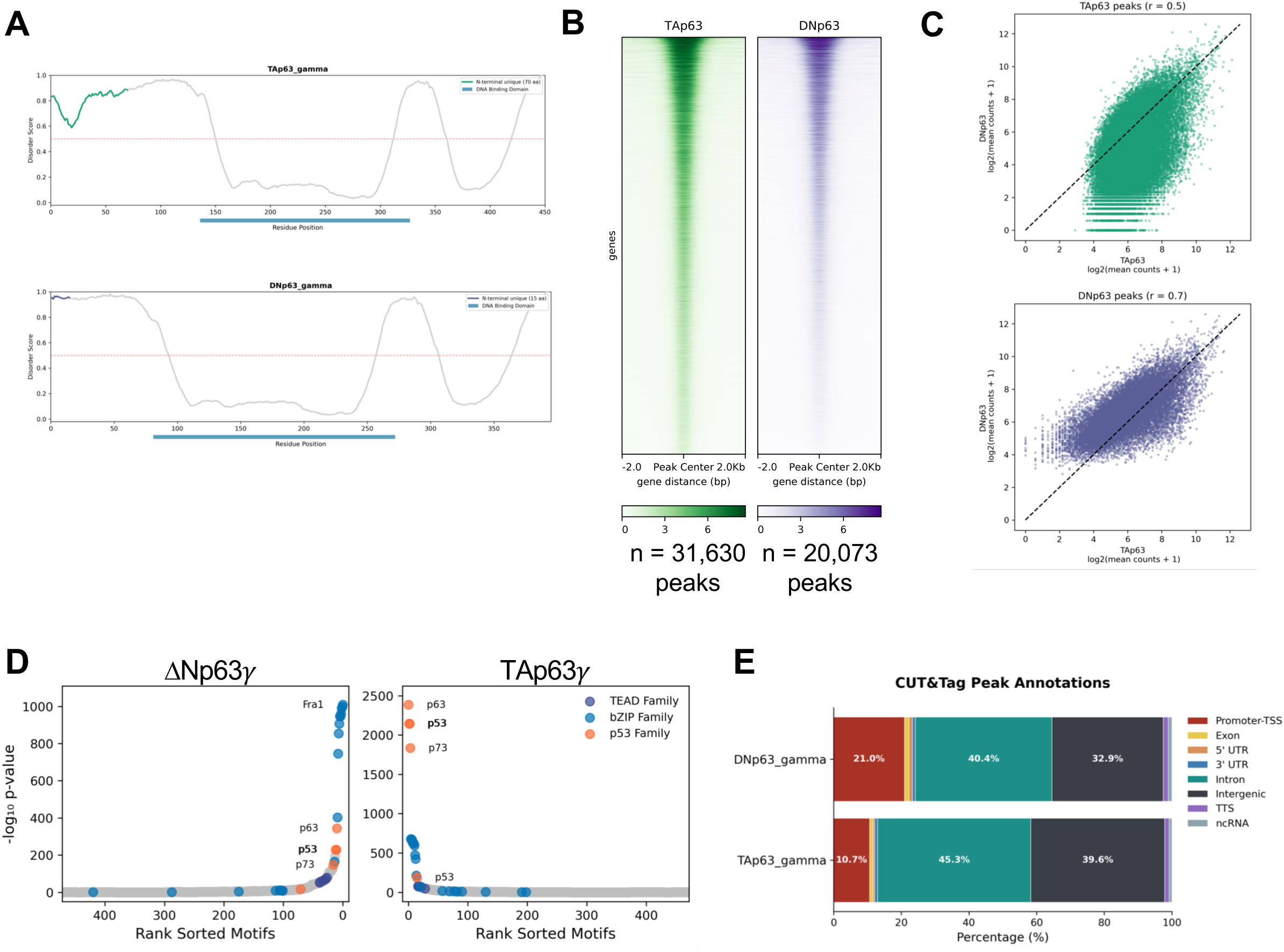
Genomic occupancy of TAp63*γ*; and ΔNp63*γ*; using CUT&Tag. **(A)** Metapredict v3 disorder scores for the TAp63*γ*; and ΔNp63*γ*; sequences. Sequences for amino acids unique to TAp63 are in green and purple for ΔNp63. The DBD is highlighted in blue at the bottom of the plot. Scores above 0.5 are predictive of disorder. **(B)** Heatmap showing TAp63*γ*; and ΔNp63*γ*; occupancy across all merged CUT&Tag peaks (n=51,703 peaks). **(C)** Correlation of TAp63*γ*; peaks between TAp63*γ*; and ΔNp63*γ*; (green), and correlation of ΔNp63*γ*; peaks between TAp63*γ*; and ΔNp63*γ*; (purple). The Pearson correlation coefficient is defined as r. **(D)** HOMER motif enrichment analysis of ΔNp63*γ*; peaks (left) and TAp63*γ*; peaks (right). **(E)** Peak annotations for TAp63*γ*; and ΔNp63*γ*; peaks.

To investigate the possibility that ΔNp63 retains transcriptional activity, we investigated the physical characteristics of the N-terminus of the p63 isoforms. ADs are composed of a balance of acidic and hydrophobic residues (51–53). Based off a simple prediction formula developed by Kothka and Staller et al. 2023 (54), we analyzed the amino acid sequence of TAp63*γ*; and ΔNp63*γ*; in segments of 39 amino acids and calculated the charge and hydrophobicity of each window (**Fig. S3A**). We also analyzed p53 as a comparison since it is known to share AD features with p63 (**Fig. S3B**). While the model predicts that ADs have a net charge ≤ −8 and a WFL count ≥ 6, we found that neither TAp63 nor ΔNp63 uniformly reached the threshold for hydrophobicity or charge in the N-terminus like p53, emphasizing that there is more complexity to determining a robust AD. However, both the N-terminus of TAp63 and ΔNp63 exhibited a net negative charge, consistent with the historical characterization of an AD as an “acidic blob” (55). These observations provide additional theoretical support for an AD on ΔNp63. In summary, analysis of the N-terminal amino acid sequences of TAp63 and ΔNp63 revealed that the isoforms share an IDR leading up to the DBD, but the N-terminus of TAp63 is characterized by a known alpha-helix, a longer IDR, and a more acidic peptide sequence.

### TAp63 and ΔNp63 broadly share genomic occupancy

Since there has been emerging evidence that IDRs can shape TF binding preferences (59, 60), we asked whether differences in binding of TAp63 and ΔNp63 could explain the distinct transcriptional effects of the isoforms. We used CUT&Tag to profile TF binding with a pan-p63 antibody and a His-tag antibody following induction of His-tagged TAp63*γ*; and ΔNp63*γ*; in PCI-30 for 24 hours. We identified 31,630 and 20,073 peaks following overexpression of TAp63*γ*; and ΔNp63*γ*;, respectively (**Fig. 2B**). Reassuringly, we observed a strong correlation between called peaks across the different antibodies (Pearson correlation coefficient r > 0.9) (**Fig. S4B**). Between the isoforms, 13,198 peaks were shared (**Fig. S4C**). We then looked at the correlation between isoforms of peaks called in either the TAp63*γ*; condition or the ΔNp63*γ*; condition (**Fig. 2C**). We observed that TAp63 and ΔNp63 display a strong positive correlation (Pearson correlation r = 0.7) for peaks called in the ΔNp63 condition and a modestly positive correlation (r = 0.5) for peaks called in the TAp63 condition. Overall, these results indicate that TAp63 and ΔNp63 broadly share genome occupancy, though there are a few peaks that are unique to TAp63, suggesting that the DBD is primarily responsible for directing p63 occupancy.

### The N-terminus of TAp63 shapes distal, intergenic binding preferences

Using k-means clustering, we identified subsets of peaks that differed primarily in signal between the isoforms (**Fig. S4A**). While clusters 2 and 3 represent peaks with stronger signal in either TAp63 or ΔNp63, only cluster 3 stood out as peaks that could be *bona fide* TAp63-specific. Though there was not robust evidence for peaks that are unique to ΔNp63, we found that TF motifs in the TAp63 condition were more enriched for the p53 family than motifs in the ΔNp63 condition (**Fig. 2D**). Surprisingly, a greater percentage of peaks bound by ΔNp63 were annotated as near the transcription start site (TSS) as compared to TAp63 (21.0% versus 10.7%) (**Fig. 2E**), indicating that TAp63 binds more regulatory elements further away from the TSS than ΔNp63. Consistent with this observation, we found that the top differentially bound peaks unique to TAp63 are located ≥ 10kb from a TSS (**Fig. S2D**). Therefore, comparing ΔNp63 binding to TAp63, ΔNp63 is characterized by increased association with proximal promoter regions enriched for activating TF motifs such as the AP-1 family, whereas TAp63 is characterized by increased binding to distal regulatory elements.

### TAp63 and ΔNp63 exhibit similar binding near the promoters of genes involved in apoptosis

Having characterized the binding of N-terminal p63 isoforms, we asked whether differential binding could explain the discrepancies in transcriptional regulation of apoptosis that we had observed. We focused specifically on peaks that were annotated to be near genes in the MSigDB HALLMARK apoptosis gene set and found that the vast majority of peaks were bound by both TAp63 and ΔNp63 (**Fig. 3A**). Specifically, peaks that were annotated to be near genes involved in apoptosis showed a strong correlation between TAp63 and ΔNp63 (Pearson correlation r = 0.7) (**Fig. 3B**). We therefore focused our attention on key drivers of apoptosis. We honed in on genes *BBC3* and *PMAIP1*, which encode for PUMA and NOXA, respectively, since transcriptional upregulation of PUMA and NOXA is known to occur in a p53-dependent manner (61, 62). We found that both TAp63 and ΔNp63 localized to the promoters of these genes (**Fig. 3C**). We observed a similar pattern at the promoter of *GPX1*, a gene also found in the apoptotic signature. These data indicate that while both TAp63 and ΔNp63 bind near or to the promoters of genes regulating apoptosis, only TAp63 is capable of transcriptionally activating these target genes.

**Figure 3.**
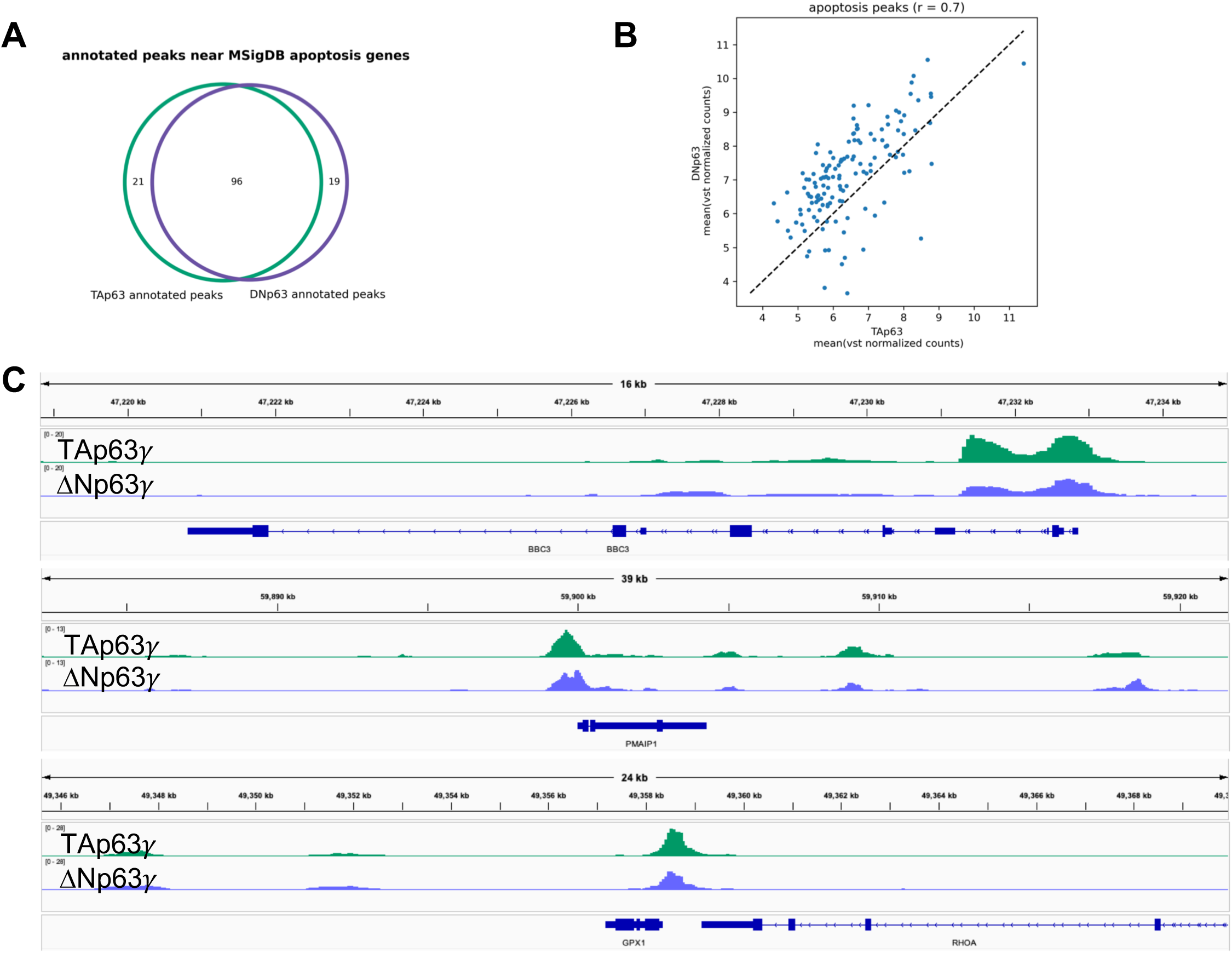
TAp63*γ*; and ΔNp63*γ*; exhibit similar binding near promoters of genes involved in apoptosis. **(A)** Overlap of annotated peaks found in the MSigDB HALLMARK apoptosis gene set. **(B)** Pearson correlation of TAp63*γ*; and ΔNp63*γ*; peaks annotated to be near genes found in the MSigDB HALLMARK apoptosis gene set. The Pearson correlation coefficient is defined as r. **(C)** Genome browser screenshots of TAp63*γ*; and ΔNp63*γ*; binding near the promoters of genes involved in apoptosis, *BBC3, PMAIP1, and GPX1*.

Naturally, these observations raised the possibility that ΔNp63 is functioning as a repressor at the genes involved in transcriptional regulation of apoptosis. Genes from the apoptosis signature were not uniformly downregulated upon expression of ΔNp63 isoforms (**Fig. 1E**). To explore the possibility that ΔNp63 functions solely as a repressor more broadly, we used our transcriptomic data and loosely defined a DEG as upregulated if the log_2_fold change (log_2_FC) > 0.6 or downregulated if the log_2_FC < −0.6. We annotated peaks with the nearest gene if the peak was located 10kb upstream or 2kb downstream of the TSS. We then intersected the upregulated and downregulated DEG for each isoform with their respective peaks to determine binding events that result in gene regulatory effects (**Fig. 4A**). We found that 42% of upregulated DEG by TAp63 and 31% of upregulated DEG by ΔNp63 were associated with TF binding near the TSS (**Fig. 4B**). On the other hand, 23% of downregulated DEG by TAp63 and 26% of downregulated DEG by ΔNp63 were associated with TF binding (**Fig. 4B**). These observations highlight that binding of TAp63 and ΔNp63 generally leads to increased gene expression rather than downregulation, and that binding of TAp63 results in more transcriptomic changes than ΔNp63. Most importantly, these observations suggest that ΔNp63 is not imbued with uniquely repressive functions in this context.

**Figure 4.**
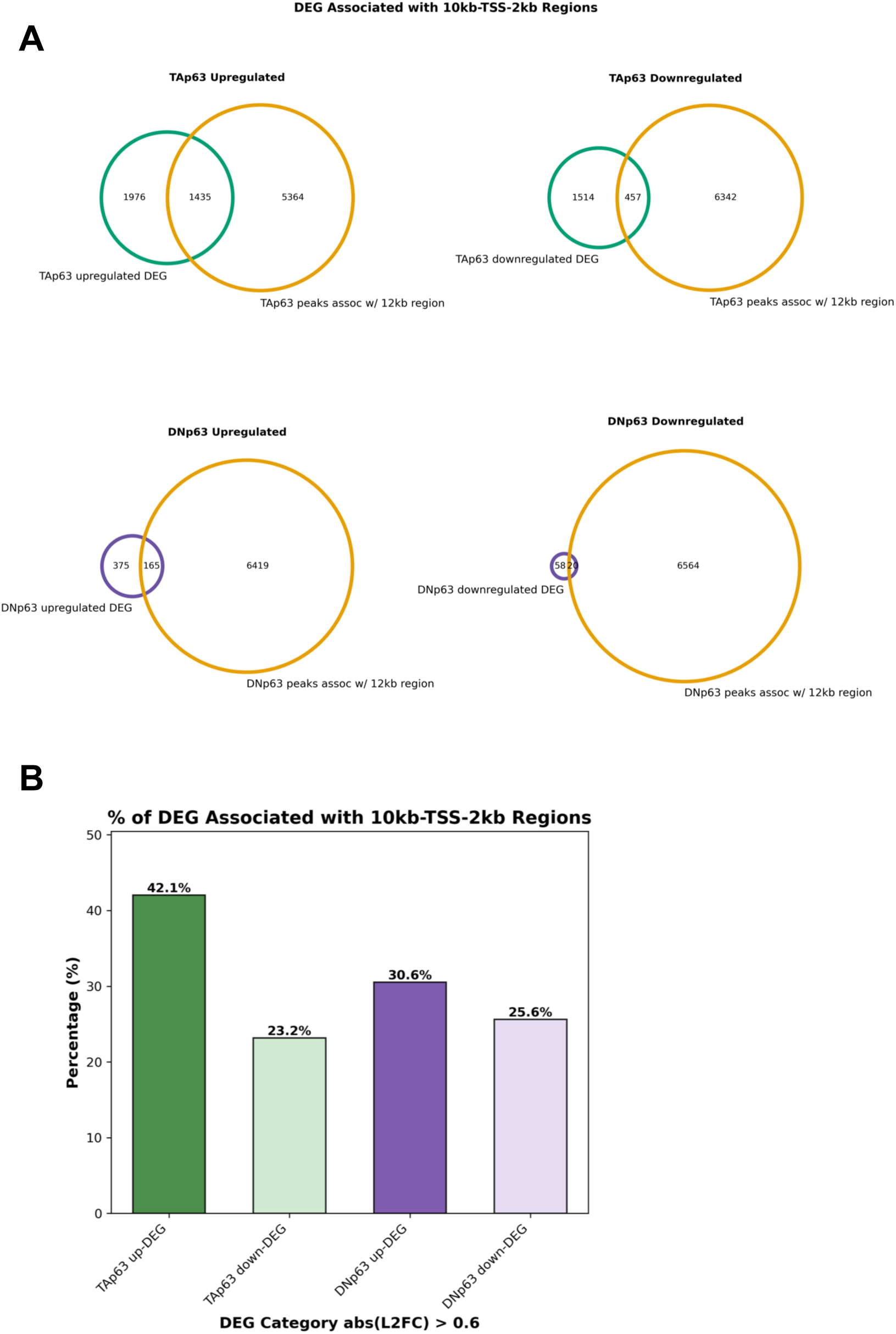
Intersection of DEGs and peaks reveals effect of binding of TAp63*γ*; and ΔNp63*γ*; on gene expression. **(A)** Overlap of peaks within 10kb upstream or 2kb downstream of a gene’s TSS and up- or down-regulated DEG following expression of TAp63*γ*; and ΔNp63*γ*;. Here, DEGs are defined as having an absolute log_2_fold change > 0.6 and an adjusted p-value < 0.01. **(B)** Percentage of DEG for each isoform that are associated with either upregulation or downregulation upon expression of the respective isoform.

### TurboID proximity labeling defines the fuzzy interactomes of p63 isoforms

Given that TAp63 and ΔNp63 broadly target similar genomic sites and bind near the promoters of pro-apoptotic genes, we considered how differences in protein-protein interactions could influence gene regulation by the p63 isoforms. IDRs in ADs are known to mediate weak and “fuzzy” interactions with other IDRs of coactivators, corepressors, and chromatin remodelers. Because of the predicted disorder of the N-terminus in the p63 isoforms (**Fig. 2A**), we used a proximity labeling approach followed by mass spectrometry (MS) to identify proteins that neighbor or weakly interact with the isoforms. We fused the biotin ligase TurboID to the N-terminus of each p63 isoform and transfected these fusion constructs into PCI-30 (**Fig. 5A, S5A, S5B**). We added a nuclear localization signal (3xNLS) to TurboID to control for nonspecific labeling in the nucleus. In total, we identified between 169-251 proteins in the interactomes for TAp63⍺, ΔNp63⍺, ΔNp63*γ*;, and TAp63*γ*; (**Fig. S5C**). Consistent with previous studies that have shown TAp63⍺ is in a closed, inactive confirmation prior to phosphorylation of the ID (19), the interactome of TAp63*γ*; was larger than TAp63⍺ (245 versus 169 enriched proteins) and encompassed most of the TAp63⍺ interactome (**Fig. S5D**). Surprisingly, we found that the *γ*; C-terminus of both TAp63 and ΔNp63 mediated over twice the number of unique protein interactions than the ⍺ C-terminus, revealing that despite losing the structured SAM domain in the ⍺ terminus, the *γ*; C-terminus is capable of its own specific interactions.

**Figure 5.**
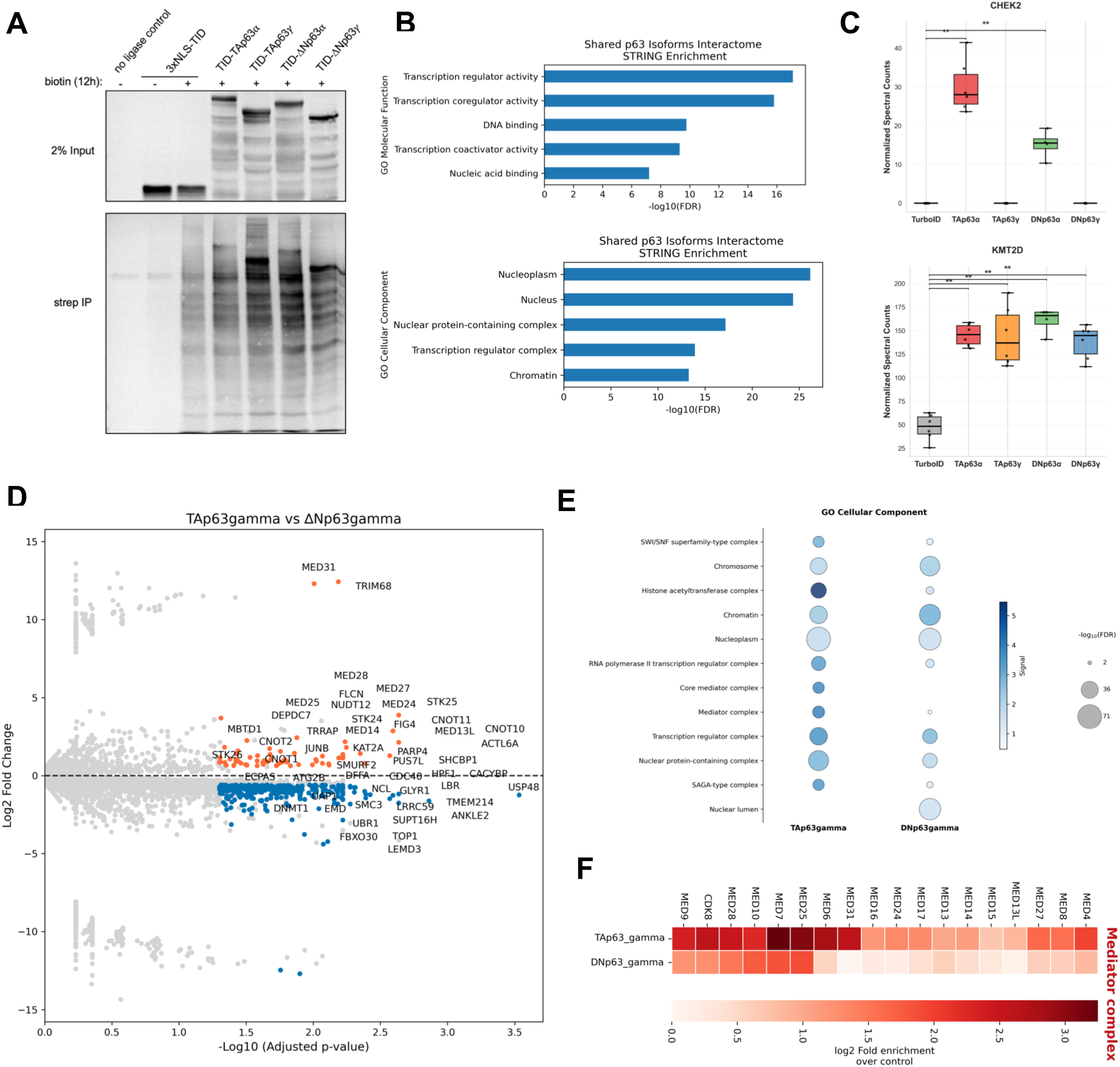
TurboID proximity labeling maps the interactomes of p63 isoforms. **(A)** Immunoblot for input and anti-streptavidin IP from PCI-30 cells expressing p63 isoforms fused to biotin ligase TurboID (TID). **(B)** Enriched GO Molecular Function and Cellular Component terms for the shared p63 isoform interactome. **(C)** Normalized MS spectral counts of TurboID biotinylated CHEK2 and KMT2D. Mann-Whitney U test comparing each isoform to TurboID alone. Each point represents one sample, bar indicates mean, box indicates 95% confidence interval, and whiskers indicate the range. n = 5-6, *p<0.05, **p<0.01, ***p<0.001. **(D)** Volcano plot of the TAp63*γ*; (orange) versus ΔNp63*γ*; (blue) interactomes. Proteins with an absolute log_2_FC > 1 and a p_adj_ < 0.05 are in color. Non-significant proteins are in gray. **(E)** Enrichment analysis of the TAp63*γ*; and ΔNp63*γ*; interactomes using GO Cellular component terms. **(F)** Heatmap of log_2_fold enrichment for subunits of the Mediator complex for the TAp63*γ*; interactome and ΔNp63*γ*; interactome.

Reassuringly, the combined interactomes for all p63 isoforms were enriched for Gene Ontology Cellular Component and Molecular Function terms related to the nucleus and transcription regulatory activity (**Fig. 5B**). Additionally, our experiment identified proteins previously known to interact with p63, including KMT2D and CHEK2 (63, 64). CHEK2 is the kinase responsible for phosphorylating and activating TAp63⍺ in conditions of stress (63). Concordantly, the ⍺ C-terminal isoforms pulled-down CHEK2, but not the *γ*; isoforms (**Fig. 5C**). Though there was an enrichment for DNA-independent cofactors in our experiment, we also identified TFs that neighbor all p63 isoforms, including Gli2 and Sox13 (**Fig. S5E**). p63 may either directly interact with these TFs through the DBD or another shared domain, or p63 may neighbor these TFs by localizing to similar regulatory elements in the genome. A full list of proteins for each isoform is available in **Supplementary Table X**. To our knowledge, this is the first isoform-specific interactome data available for p63.

### The TAp63 interactome is enriched for transcriptional coactivators and histone acetyltransferases

To determine how differences in the N-terminus of TAp63 and ΔNp63 could influence interactions with other proteins, we specifically compared the interactomes of TAp63*γ*; and ΔNp63*γ*; (**Fig. 5D**). Through STRING network analysis of the interactomes, it was immediately apparent that TAp63*γ*; neighbors more coactivator complexes than ΔNp63*γ*; (**Fig. S6A and S6B**). Enrichment analysis showed that compared to ΔNp63*γ*;, the interactome of TAp63*γ*; is enriched for cofactors necessary for transcriptional activation, and specifically components of the SWI/SNF, histone acetyltransferase, Mediator, and SAGA complexes (**Fig. 5E**). More specifically, the interactome of TAp63*γ*; is enriched for several subunits of the transcriptional coactivator Mediator (**Fig. 5F**). Critically, we observed that ΔNp63*γ*; neighbors many of these cofactors, but displays less enrichment for them than TAp63*γ*;.

Despite the inability of ΔNp63 to transcriptionally regulate apoptosis, we did not observe any enrichment in the ΔNp63*γ*; interactome for transcriptional repressor complexes or histone deacetylases (**Fig. S6C**), suggesting that ΔNp63 is not imbued with a strong repressive function. Instead, both TAp63 and ΔNp63 neighbored proteins that can be classified as negative regulators of transcription using GO Biological Process terms (**Fig. S6D**). Since previous studies have shown that ΔNp63 interacts with the repressive histone deacetylases HDAC1 and HDAC2 (65, 66), we specifically looked for these proteins in our dataset. However, none of the p63 isoforms showed increased pull-down for HDAC1 or HDAC2 compared to the control in our cell line (**Fig. S6E**). We did observe enrichment of the histone deacetylases HDAC4 and HDAC7 with all isoforms of p63 (**Fig. S6E**). Together with the transcriptomic and binding data, these results emphasize that ΔNp63 does not preferentially interact with transcriptional repressors as compared to TAp63. Instead, while both TAp63 and ΔNp63 neighbor coactivators, the interactome of TAp63 is heavily enriched for proteins critical in transcriptional activation, specifically subunits of mediator, histone acetyltransferases, and the SAGA complex.

### Cofactor interactions with TAp63 influence chromatin accessibility and TF binding at intergenic sites

Given that the interactome of TAp63 was enriched for components of the SWI/SNF complex, histone acetyltransferases, and the SAGA complex, we then asked if TAp63 and ΔNp63 have different effects on chromatin accessibility. Using ATAC-sequencing, we profiled chromatin accessibility following doxycycline induction of TAp63*γ*; and ΔNp63*γ*; in PCI-30 for 24 hours. We defined differentially accessible regions (DARs) as peaks that displayed increased accessibility over a GFP-expressing control line also induced with doxycycline. Using principle component analysis (PCA), we noted that ΔNp63*γ*; clustered near the control samples (**Fig. 6A**). Analyzing the DARs, expression of ΔNp63*γ*; resulted in very few changes in chromatin accessibility (**Fig. 6B**), while overexpression of TAp63*γ*; led to over 3,000 regions with increased accessibility (**Fig. 6C**). These regions were enriched for motifs in the p53 family, suggesting that the increased accessibility is a direct result of TAp63 binding. We then hypothesized that these DARs could have an effect on gene regulation at or near the promoter. Intersecting the DARs with upregulated DEG from the transcriptome data, we found few examples of increased accessibility near the promoter of genes upregulated by TAp63 (**Fig. S7A**). Most genes, including genes involved in apoptosis, exhibited accessible regions near the promoter in the control condition, revealing that for many of the genes regulated by the p63 isoforms, TAp63 and ΔNp63 are binding already accessible loci (**Fig. 6E**). By contrast, we found that the majority of DARs upon TAp63 expression are located in intronic or intergenic regions (**Fig. 6D**). This observation was consistent with our earlier finding that TAp63 binds more distal regulatory elements than ΔNp63 (**Fig. 2E**). We found that many of the most significant DARs were located at the same sites we found unique to TAp63 binding from the CUT&Tag data (**Fig. 6F and S6B**). Collectively, our observations indicate that unique binding by TAp63 at distal or intergenic sites can be attributed to an increase in accessibility upon TAp63 expression. Together with the TurboID interactome data, we hypothesize that TAp63 recruits histone acetyltransferases and chromatin remodelers through its unique N-terminal region to bind DNA at previously inaccessible sites.

**Figure 6.**
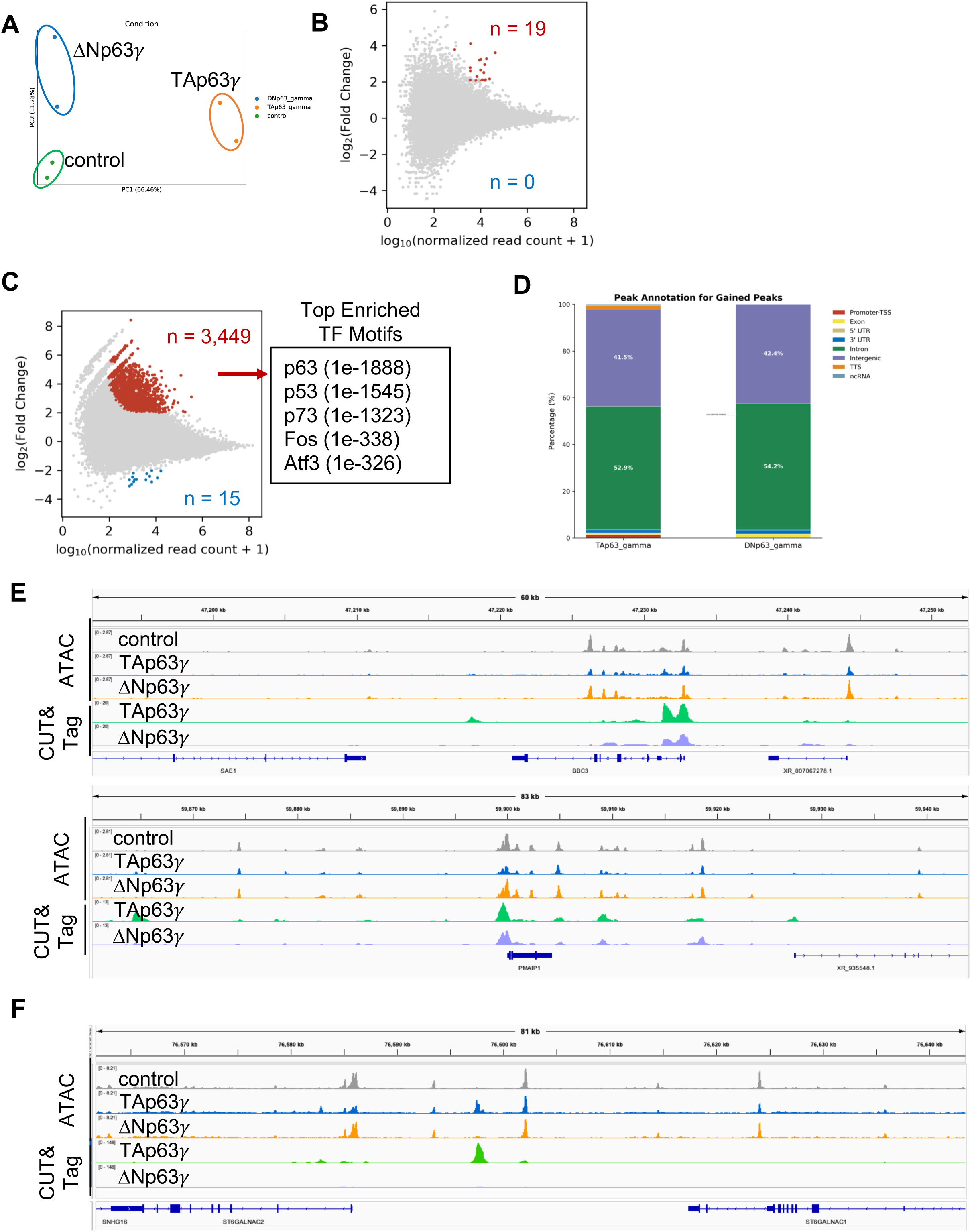
Chromatin accessibility following expression of TAp63*γ*; and ΔNp63*γ*;. **(A)** Principle component analysis of normalized counts from ATAC-seq in cell lines induced with doxycycline to express either GFP (control), TAp63*γ*;, or ΔNp63*γ*;. **(B)** Differentially accessible regions (DARs) following expression of ΔNp63*γ*; compared to control. **(C)** DARs following expression of ΔTAp63*γ*; compared to control and the top enriched motifs found in the regions displaying increased accessibility. **(D)** Annotations for regions that gained accessibility following expression of TAp63*γ*; or ΔNp63*γ*;. **(E)** Genome browser screenshots of CUT&Tag peaks and chromatin accessibility (ATAC-seq) following expression of TAp63*γ*; and ΔNp63*γ*;.

### Predicted IDR of TAp63 mediates interaction with KAT2A

From the interactome data, one of the most enriched proteins found to be neighboring TAp63 was the histone acetyltransferase KAT2A, a member of the Spt-Ada-Gcn5 acetyltransferase (SAGA) complex which is important for transcriptional activation (67–69). We compared TAp63 and ΔNp63 interactions with the cryo-EM structure of human SAGA complex (68) and found that the TAp63 interactome exhibits increased enrichment for KAT2A, SGF29, TRRAP, TAF12, and ATXN7 (**Fig. 7A**). Notably, KAT2A and SGF29 are two of four proteins in the histone acetyltransferase (HAT) module in SAGA, the other two components are TADA2B and TADA3 which displayed slight increases in enrichment. ATXN7 is part of the deubiquitinase (DUB) module, which interfaces with the HAT module in the cryo-EM structure. We did not observe significant enrichment with components of the scaffolding core module that includes TATA-box binding protein (TBP)-associated factors (TAFs). TurboID labels a cloud of proteins nearby, thus these observations indicate that there is a degree of spatial specificity that can be achieved with TurboID labeling.

**Figure 7.**
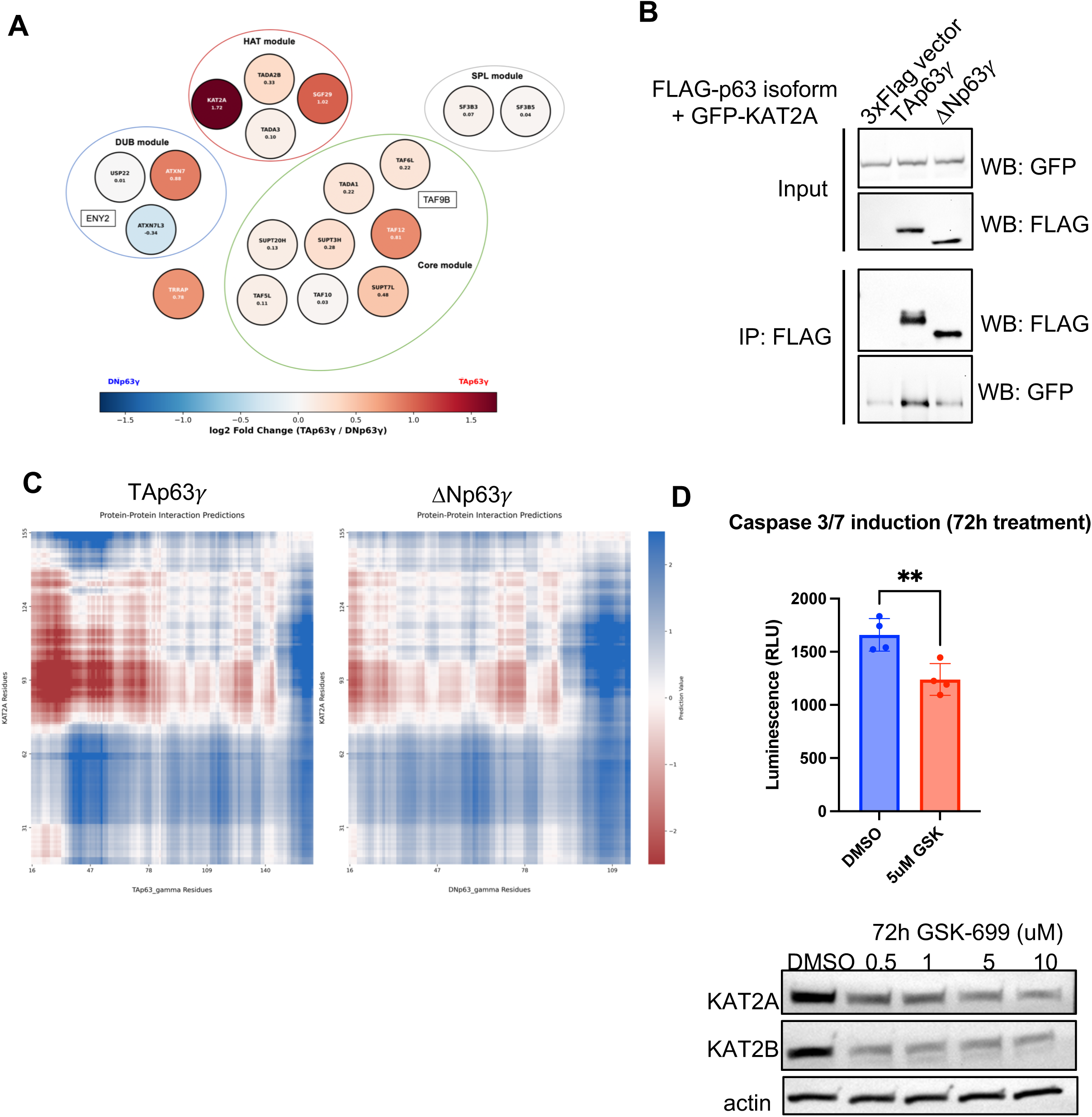
TAp63 cooperates with KAT2A to transcriptionally regulate apoptosis. **(A)** Diagram of members of the SAGA complex that were identified in the TurboID-MS dataset. Colors indicate enrichment in the TAp63*γ*; interactome compared to the ΔNp63*γ*; interactome. The log_2_FC of normalized spectral counts is written inside each circle. **(B)** Coimmunoprecipitation of FLAG-tagged p63 isoforms with GFP-tagged KAT2A. **(C)** FINCHES protein-protein interaction predictions between TAp63*γ*; or ΔNp63*γ*; and KAT2A. **(D)** Luminescence values of caspase 3/7 activation in PCI-30 cells induced to express TAp63*γ*; and treated with 5uM GSK-699 for 72 hours. Immunoblot of KAT2A and KAT2B following different concentrations of GSK-699 below.

To validate the enriched interaction between TAp63 andKAT2A, we first performed a co-immunoprecipitation in 293T using exogenous GFP-tagged KAT2A with either FLAG-tagged TAp63*γ*; or ΔNp63*γ*;. We validated that TAp63*γ*; pulled-down more KAT2A than ΔNp63*γ*; (**Fig. 7B**), indicating that TAp63 preferentially interacts with KAT2A. Since the N-terminus of TAp63 differs from ΔNp63 in predicted length of the IDR, we used FINCHES to predict interactions between disordered regions of the p63 isoforms and KAT2A (42, 43). We found that the IDR of TAp63 is predicted to more strongly interact with the IDR of KAT2A (**Fig. 7C**). These predictions suggest that the increased IDR in the N-terminus of TAp63 mediates stronger cofactor interactions than the N-terminus of ΔNp63. Our data serves as a proof of concept that a proximity-based labeling approach can identify weak and fuzzy interactions likely mediated by IDRs that would be challenging to identify through a traditional IP-MS approach that requires more stable interactions, while simultaneously uncovering isoform-specific interactions and biology as it pertains to p63 specifically.

### TAp63 cooperates with KAT2A to transcriptionally regulate apoptosis

Given the isoform-specific interaction enrichment between KAT2A and TAp63*γ*; but not ΔNp63*γ*;, we hypothesized that KAT2A may preferentially interact with TAp63 at promoters critical for the regulation of apoptosis. We therefore first treated PCI-30 cells expressing a doxycycline-inducible TAp63*γ*; with the PROTAC GSK-699 (70), which degrades KAT2A and its paralogue KAT2B, and then measured caspase 3/7 activation with a luminescence assay (**Fig. 7D**). KAT2B can be interchangeable with KAT2A in the SAGA complex in a mutually exclusive manner (71–73). We found a modest but significant decrease in TAp63*γ*;-induced apoptosis upon treatment with GSK-699, indicating that KAT2A and KAT2B may be important in the transcriptional activation of pro-apoptotic genes.

## DISCUSSION

The N-terminal p63 isoforms TAp63 and ΔNp63 have long been established as having unique roles in development and cancer, but how isoform-specific regulation is encoded by the N-terminus has not been fully elucidated. Using the regulation of apoptosis as a case study and the C-terminal *γ*; variants of p63, our study provides mechanistic insight into how TAp63 and ΔNp63 can achieve distinct gene regulatory effects. Here, we provide evidence that TAp63 and ΔNp63 are both predicted to have a disordered N-terminus. We show that they broadly bind to similar sites across the genome, and that differences in protein-protein interactions with chromatin modifiers and transcriptional coactivators can explain unique binding events and influence the transactivation potential of TAp63. Specifically, we demonstrate that TAp63 cooperates with the histone acetyltransferase KAT2A to regulate apoptosis.

Strikingly, the amino acid sequences of TAp63 and ΔNp63 are remarkably similar – there are only 15 amino acids unique to the ΔNp63 N-terminus, and 70 amino acids unique to TAp63. The widely posited explanations of the divergent functions of the isoforms are that ΔNp63 is repressive, incapable of transactivation, and/or functions as a dominant negative to inhibit activity of TAp63. However, we did not find substantial evidence that ΔNp63 lacks transactivation potential or that it is imbued with uniquely repressive functions not found in TAp63. First, we found that the N-terminus of ΔNp63 is acidic and that it is predicted to be disordered – both widely accepted features of activation domains. Second, motifs found in peaks occupied by ΔNp63 were enriched for activating TFs like the AP-1 family. Third, we found that binding of TAp63 and ΔNp63 resulted in similar rates of downregulation of nearby target genes and that binding of ΔNp63 also resulted in upregulation of genes. Lastly, the interactome data showed no enrichment for proteins associated with repression in the interactome of ΔNp63. Both TAp63 and ΔNp63 similarly neighbored the histone deacetylases HDAC4 and HDAC7, indicating that both isoforms can have repressive functions. Additionally, ΔNp63 neighbors CBP/p300, which is a complex critical for transcriptional activation. Instead, we propose a simpler explanation – ΔNp63 is merely a weak transcriptional activator. The literature broadly supports an activating function to ΔNp63, including transactivation potential in reporter assays and direct interaction with MED12 (15, 16, 50, 74). In our data, when comparing the interactomes of TAp63 and ΔNp63, TAp63 neighbors more transcriptional machinery. We observed increased interactions with histone acetyltransferases like KAT2A and subunits of the transcriptional coactivator Mediator. Concordantly, in our transcriptomic data, expression of TAp63 resulted in over an 8-fold increase in upregulated genes compared to ΔNp63 regulation. A recent study analyzing p63-bound cis-regulatory elements also found that occupancy of ΔNp63⍺ is only weakly correlated to transcriptional output. Simply put, TAp63 appears to be a stronger transcriptional activator than ΔNp63.

The importance of intrinsically disordered regions (IDRs) in transcriptional regulation has been expanding. Studies have shown that IDR-mediated interactions between cofactors plays a role in transcriptional regulation and that IDRs may shape TF binding preferences (56, 58, 60, 75, 76). Activation domains are typically composed of IDRs, and our study found that both the N-termini of TAp63 and ΔNp63 are predicted to be disordered, with the key difference being that the length of the IDR in TAp63 is longer than ΔNp63. Given that IDRs are thought to mediate “fuzzy” interactions, our proximity-based labeling approach with TurboID followed by mass spectrometry was advantageous for identifying proteins that weakly interact with p63 or are found in the “neighborhood” of p63 isoforms. Ultimately, this approach allowed us to define a significantly larger interactome than previous IP-mass spec experiments (64). Of note, we show that the interaction between KAT2A and TAp63 is predicted to be mediated through IDRs. We hypothesize that the increase in disorder in TAp63 is responsible for the enriched interaction in comparison to ΔNp63, but future studies will need to utilize nuanced structure/function experiments to understand this more thoroughly.

IDRs have also been shown to influence TF target search and shape binding preferences beyond the DBD (58, 60). While our study broadly demonstrated that TAp63 and ΔNp63 bind similarly throughout the genome, we did observe binding that was unique to TAp63. In combination with our results showing that TAp63 expression is associated with an increase in chromatin accessibility, we found that these unique binding sites were typically at previously inaccessible regions. In addition to the findings that these accessible regions are enriched for the p63 motif and that TAp63 interacts with chromatin modifiers, we hypothesize that much of TAp63-specific binding can be attributed to its ability to recruit cofactors that relax the chromatin and allow for TAp63 binding. Contrary to some studies, we do not think that TAp63 itself has pioneering ability. Rather, it appears to rely on IDR-mediated interactions with cofactors to evict or acetylate nucleosomes and permit TF binding. KAT2A is part of the histone acetyltransferase module in the SAGA complex. Historically, TRRAP has been thought to be responsible for mediating interaction of a TF with SAGA (77, 78). However, a recent study has shown that the SAGA HAT module, which includes KAT2A, can bind both TFs and nucleosomes (78). Furthermore, cryo-EM structure of the human SAGA complex demonstrated notable differences to the yeast structure, including the lack of a subunit for TATA-box binding protein, suggesting that historical characterizations of SAGA based off the yeast complex may be different in human with functional consequences (68, 79–81). For this study, we propose a model in which KAT2A can acetylate histones to open chromatin at inaccessible sites to p63 and then facilitate TAp63 binding at its cognate motifs.

While the nucleosome eviction model with KAT2A can explain the unique TAp63 binding sites at distal, inaccessible sites, it does not account for the regulation of genes involved in apoptosis. We found that both TAp63*γ*; and ΔNp63*γ*; bind at or near the promoter of these genes and that these sites were accessible at baseline. Importantly, ΔNp63*γ*; had no effect on chromatin accessibility, underscoring that ΔNp63*γ*; cannot bind inaccessible loci like TAp63*γ*;. Since KAT2A was critical for TAp63*γ*;-induced apoptosis, we propose that the established roles of the SAGA complex in histone acetylation and transcriptional activation are dominant at these promoters. In other words, ΔNp63 cannot recruit the cofactors needed for gene activation at the response elements (RE) of apoptosis-related genes. However, this model does not account for genes that are transcriptionally activated by ΔNp63, raising the possibility that there is a difference between REs bound by ΔNp63 and are subsequently activated and those REs that are bound but not activated. This distinction could be explained by a combination of cofactor specificity at these REs and intrinsic differences in the REs for apoptosis-related genes that contain a p53 motif. A recent study using a massively parallel reporter assay (MPRA) with p63 REs showed that REs containing both the p53 and p63 motifs were significantly more active than those with a unique p63 motif, and that unique p63 RE have a higher GC content (16).

There is conflicting evidence for cofactor specificity at specific regulatory elements (75, 82–85), but our data supports cofactor specificity at p53 RE since cooperation with KAT2A at promoters was important for transcriptional regulation of apoptosis. In support of cofactor specificity, other studies have shown with MPRAs that p53-repsonsive enhancers do not require Mediator for transactivation (82). Additionally, studies in yeast suggested that housekeeping genes are mostly regulated by the general transcription factor TFIID while stress regulated genes are SAGA dependent (86–89). Together, these data support a specific role for KAT2A in transcriptional activation of apoptosis-related genes.

Collectively, our findings underscore the importance of studying individual isoforms in isolation. The isoforms of p63 add additional layers of complexity to gene regulation, and it is critical to dissect the individual contributions of each domain to understand gene regulation. Our approach serves as a blueprint for future studies of human TFs to parse out these nuanced but important differences between TF isoforms.

### Limitations of the study

Our study focused on the p63 isoforms in isolation and in a cell line lacking expression of proteins from the p53 family. Expression of ΔNp63 is predominantly in the basal epithelium while TAp63 is expressed in oocytes (90–92). Due to technical limitations, where and when TAp63 and ΔNp63 are simultaneously expressed is not well-established. There are many studies that provide evidence for broad interactions between the p53 family members, including proteomic and regulatory. To comprehensively understand the isoform-specific roles of p63, further work is needed in a tissue-specific context and with other p53 family members to determine how they function together in concert. In addition, we acknowledge that overexpression of the TFs poses its own interpretive limitations and does not necessarily model endogenous or *in vivo* behavior.

## ACKNOWLEDGEMENTS

This work was supported by R01DE032371-04 (NIH/NIDCR) (S.V.P), R01DE032865-04 (NIH/NIDCR) (S.V.P) and T32 HG000045 (NIH/NHGRI) (M.F.N). We are grateful to the GTAC at the McDonnell Genome Institute for sequencing services. For proteomic experiments, the expert technical assistance of Dr. Petra Erdmann Gilmore, Yiling Mi, Alan Davis and Rose Connors is gratefully acknowledged. The proteomic experiments were performed at the Washington University Proteomics Shared Resource (WU-PSR), R Reid Townsend MD.PhD., Director. The WU-PSR is supported in part by the WU Institute of Clinical and Translational Sciences (NCATS UL1 TR000448), the Mass Spectrometry Research Resource (NIGMS P41 GM103422; R24GM136766) and the Siteman Comprehensive Cancer Center Support Grant (NCI P30 CA091842).

## Author contributions

Marina F. Nogueira (conceptualization, data curation, formal analysis, investigation, methodology, project administration, resources, software, validation, visualization, writing-original draft), Michael J. Moore (data curation, investigation, resources, software, writing-review & editing), Aparna R. Biswas (investigation, validation), Michael P. Meers (data curation, methodology, resources, writing-review & editing), Sidharth V. Puram (conceptualization, funding acquisition, methodology, project administration, resources, supervision, writing-review & editing).

---

**Supplementary Figure S1.**
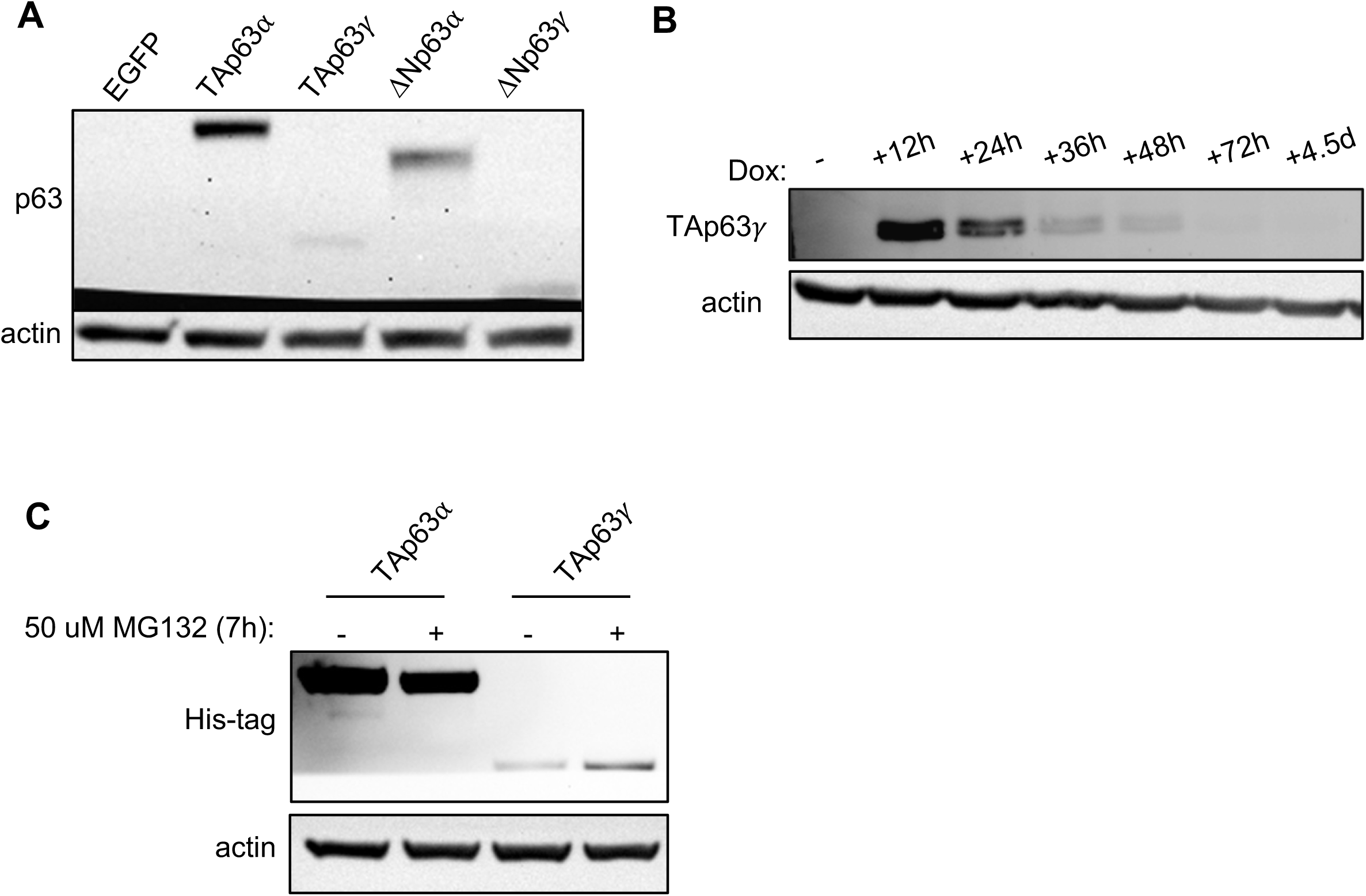
**(A)** Doxycycline (dox) induction to express p63 isoforms in PCI-30. Dox concentrations were titrated to express similar protein levels. Concentrations used were 0.5ug/mL (EGFP), 50ng/mL (TAp63⍺), 0.5ug/mL (TAp63*γ*;), 0.5ug/mL (ΔNp63⍺), 175 ng/mL (ΔNp63*γ*;). **(B)** Expression of TAp63*γ*; following dox induction. Cells were collected starting at 12 hours following dox induction up until 4.5 days. **(C)** PCI-30 cells were induced with dox for 30 hours to express each p63 isoform and treated for the last 7 hours with 50uM MG132.

**Supplementary Figure S2.**
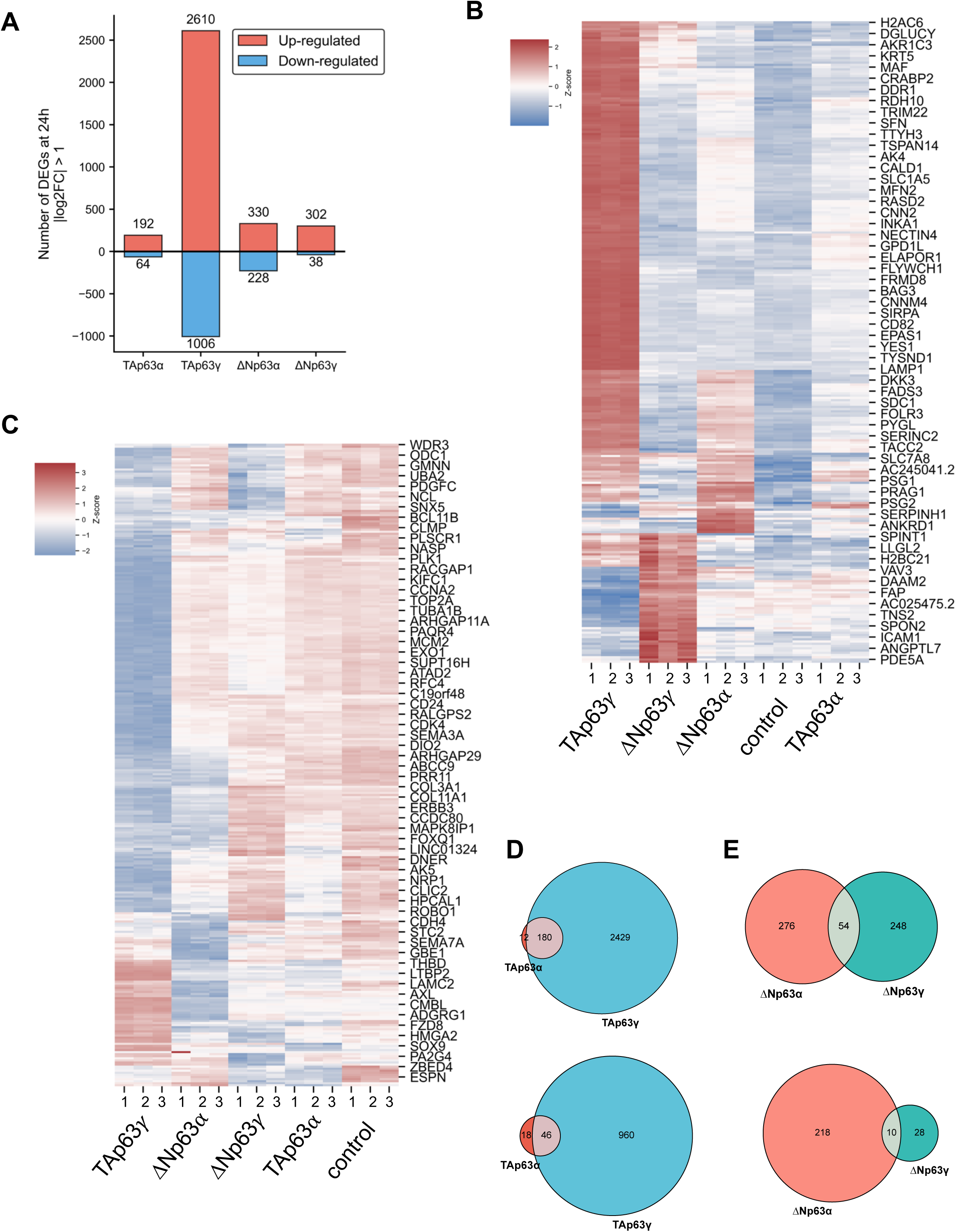
**(A)** RNA-seq following expression of each isoform. Barplots indicating the number of differentially expressed genes (DEGs) for each isoform that met an absolute log_2_FC >1 and p_adj_ < 0.05 threshold and are either up- or down-regulated in each condition. **(B)** Top 100 upregulated DEG at 24h following expression of p63 isoforms. DEG classified as log_2_FC >1 and p_adj_ < 0.05. **(C)** Top 100 downregulated DEG at 24h following expression of p63 isoforms. DEG classified as log_2_FC <1 and p_adj_ < 0.05. **(D)** Overlap of upregulated DEG by TAp63 isoforms (top) and downregulated DEG (bottom). **(E)** Overlap of upregulated DEG by ΔNp63 isoforms (top) and downregulated DEG (bottom).

**Supplementary Figure S3.**
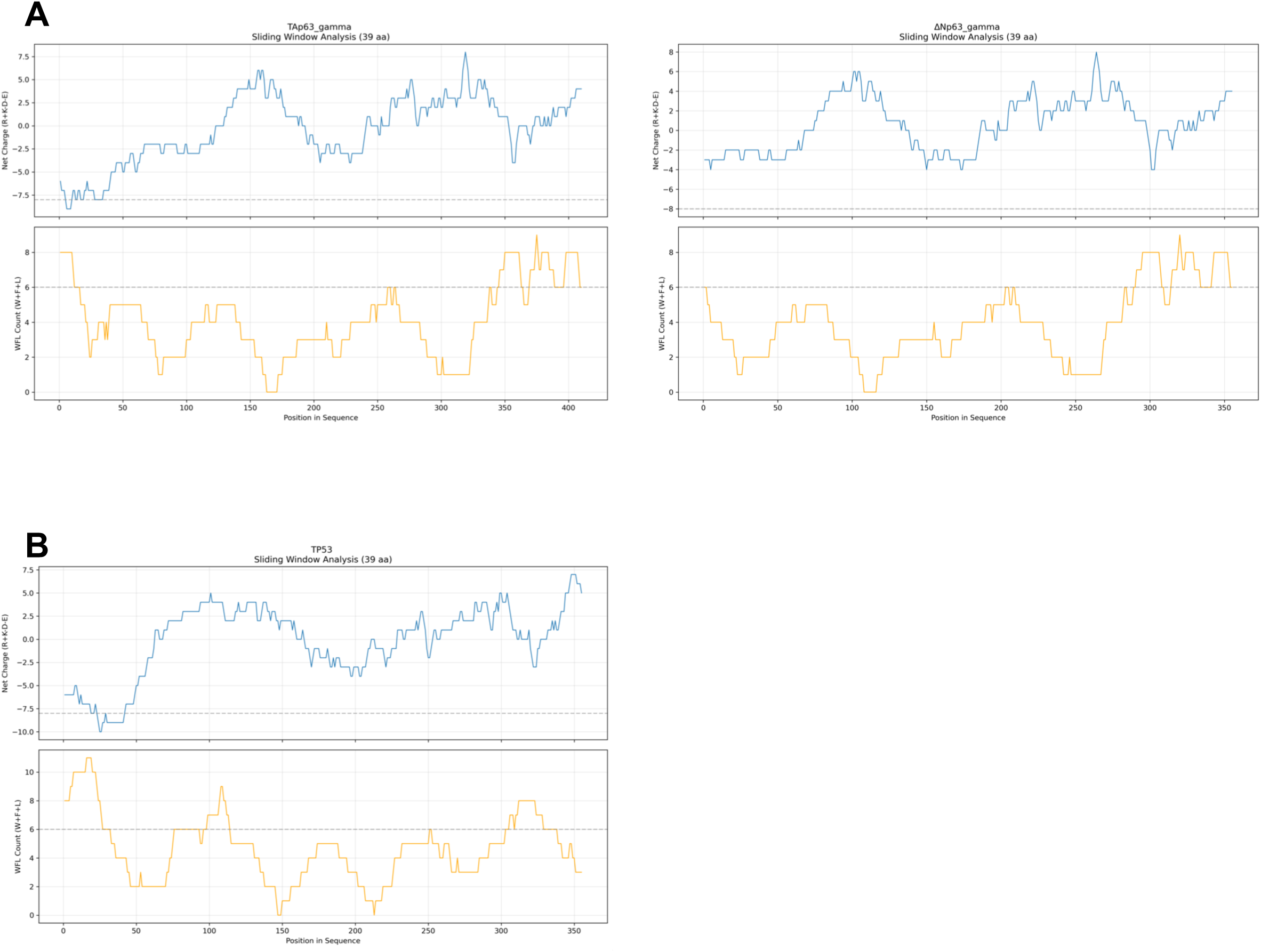
**(A)** Calculation of acidity and hydrophobicity of TAp63*γ*; and ΔNp63*γ*; using a sliding window of 39 amino acids. The net charge of a window is computed as R + K – D – E and the hydrophobicity is calculated by adding the number of W, F, and L amino acids. Activation domains are predicted to have a −13 ≥ net charge ≤ −8, and a hydrophobic count ≥ 6. Thresholds for AD prediction are indicated by the dotted line. **(B)** Calculation of acidity and hydrophobicity of p53.

**Supplementary Figure S4.**
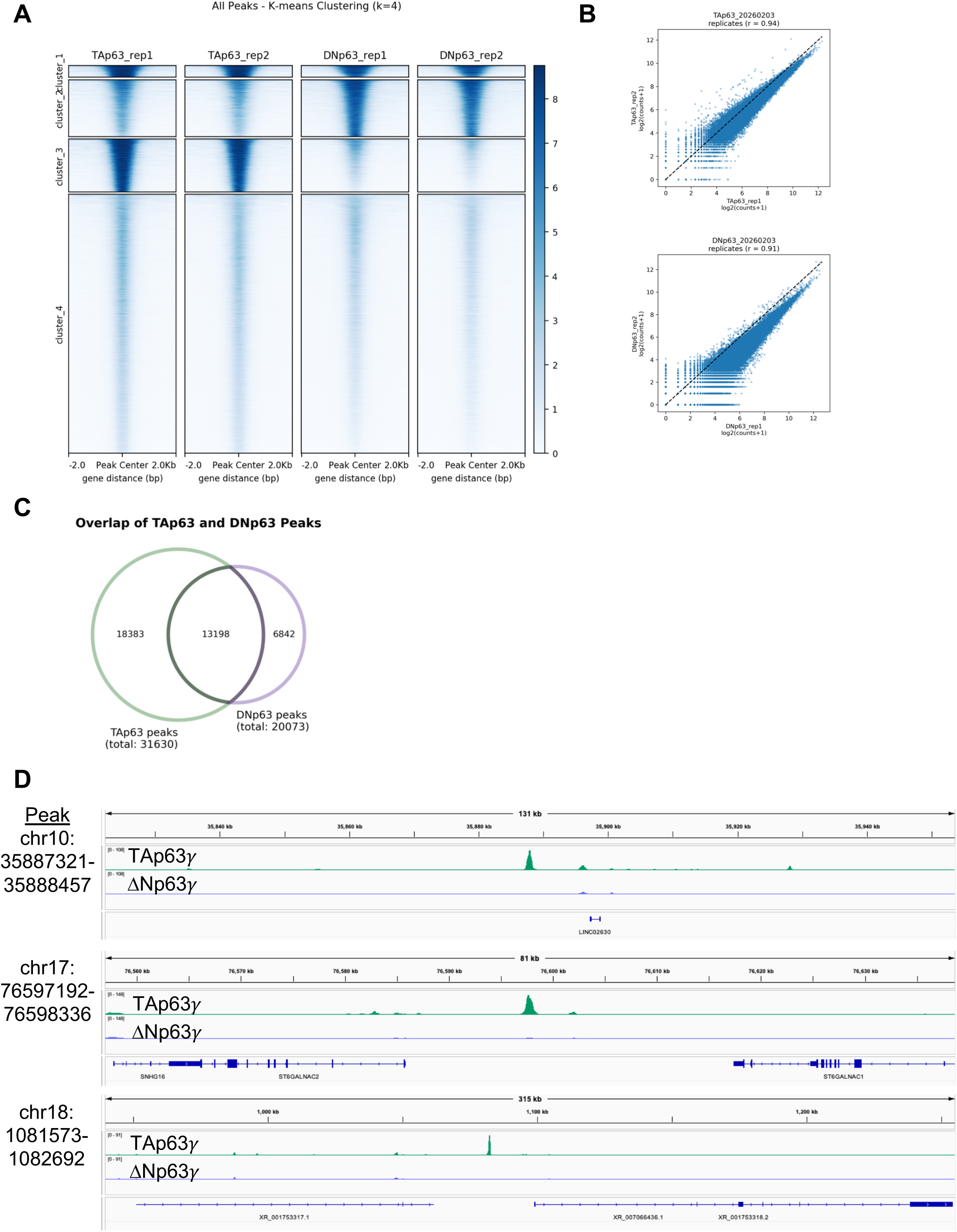
**(A)** Biological replicates of genome occupancy of TAp63*γ*; and ΔNp63*γ*; divided into 4 clusters using k-means clustering. **(B)** Correlation of MACS2-called CUT&Tag peak counts between biological replicates of TAp63*γ*; and ΔNp63*γ*;. The Pearson correlation coefficient is defined as r. **(C)** Overlap of TAp63*γ*; and ΔNp63*γ*; MACS2-called CUT&Tag peaks. **(D)** Genome browser screenshots of top differentially bound TAp63*γ*;-specific peaks located farther than 10kb from a TSS.

**Supplementary Figure S5.**
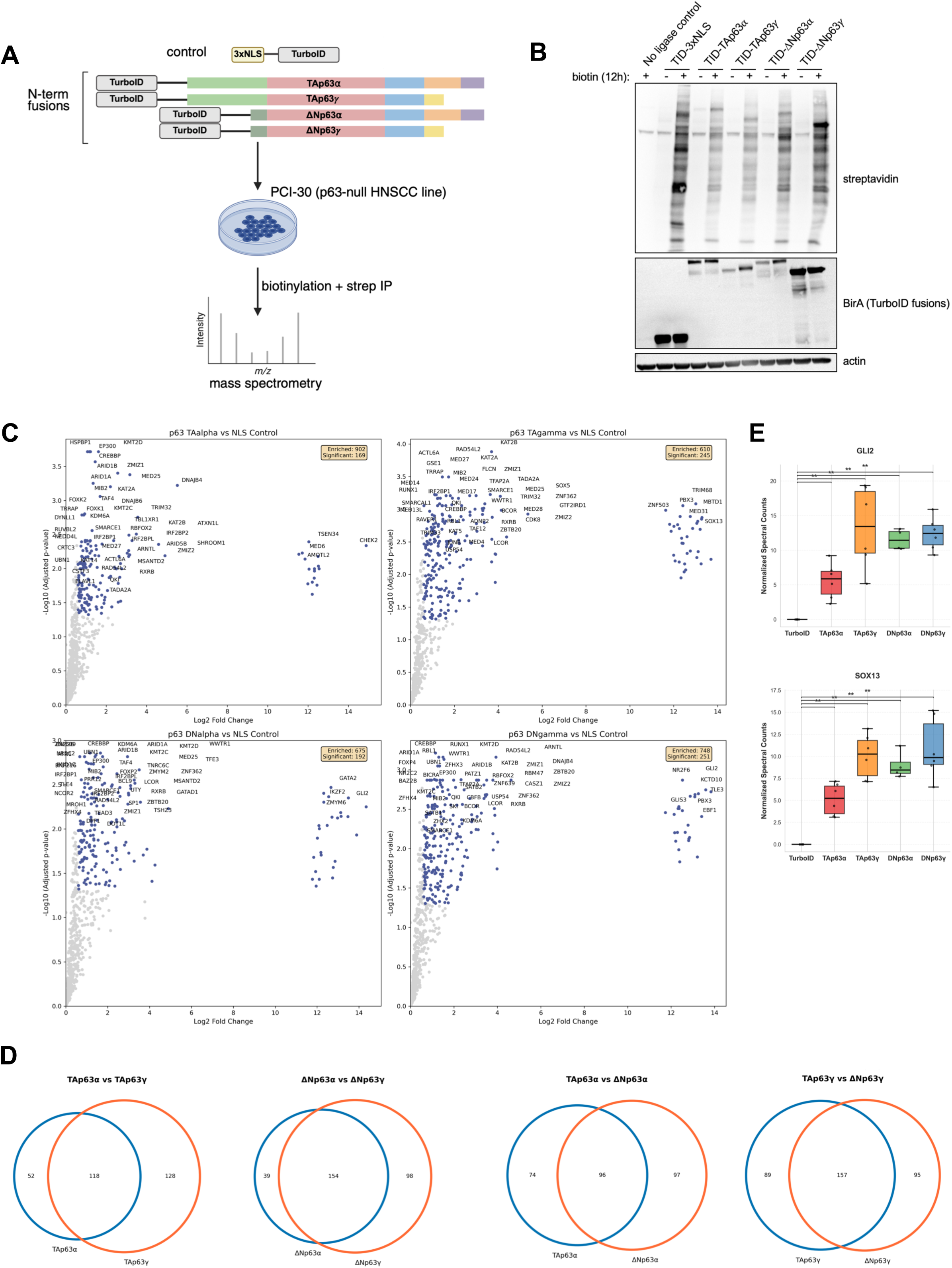
**(A)** Schematic of TurboID-mass spec (MS) labeling. **(B)** Immunoblot using whole cell lysates from PCI-30 cells expressing TurboID fused to p63 isoforms labeled with 50uM biotin for 12 hours. **(C)** Volcano plots showing enriched proteins identified by MS for each TurboID-p63 isoform compared to the 3xNLS TurboID control. **(D)** Overlap between all p63 TurboID-MS interactomes. **(E)** Normalized MS spectral counts of TurboID biotinylated Gli2 and Sox13. Mann-Whitney U test comparing each isoform to TurboID alone. Each point represents one sample, bar indicates mean, box indicates 95% confidence interval, and whiskers indicate the range. n = 5-6, *p<0.05, **p<0.01, ***p<0.001.

**Supplementary Figure S6.**
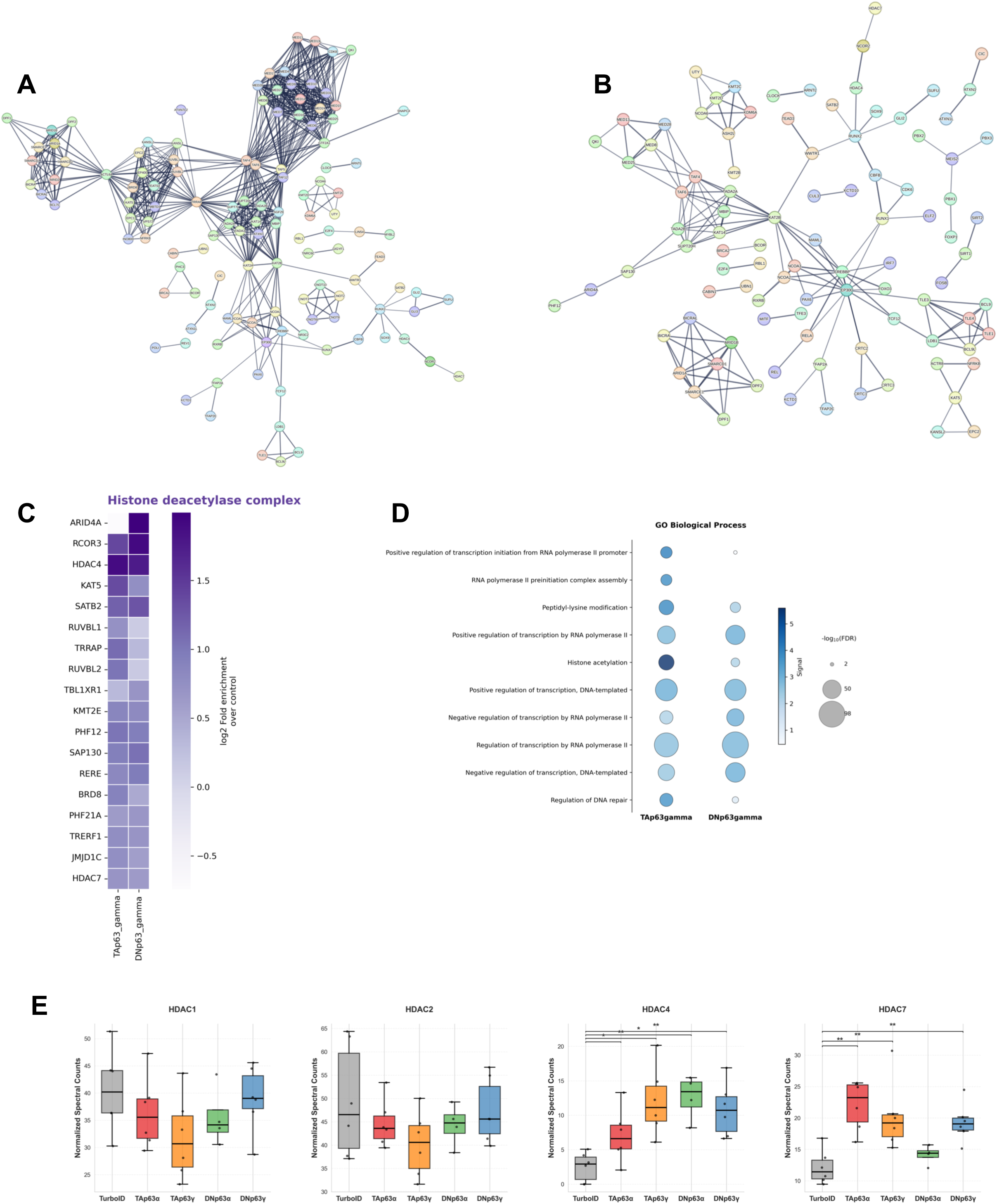
**(A)** STRING network of TAp63*γ*; interactome displaying known physical interactions. Lines indicate confidence scores > 0.7. **(B)** STRING network of ΔNp63*γ*; interactome displaying known physical interactions. Lines indicate confidence scores > 0.7. **(C)** Heatmap of log_2_fold enrichment for proteins involved in histone deacetylation for the TAp63*γ*; interactome and ΔNp63*γ*; interactome. **(D)** Enrichment analysis of the TAp63*γ*; and ΔNp63*γ*; interactomes using GO Biological Process terms. **(E)** Normalized MS spectral counts of TurboID biotinylated HDAC proteins. Mann-Whitney U test comparing each isoform to TurboID alone. Each point represents one sample, bar indicates mean, box indicates 95% confidence interval, and whiskers indicate the range. n = 5-6, *p<0.05, **p<0.01, ***p<0.001.

**Supplementary Figure S7.**
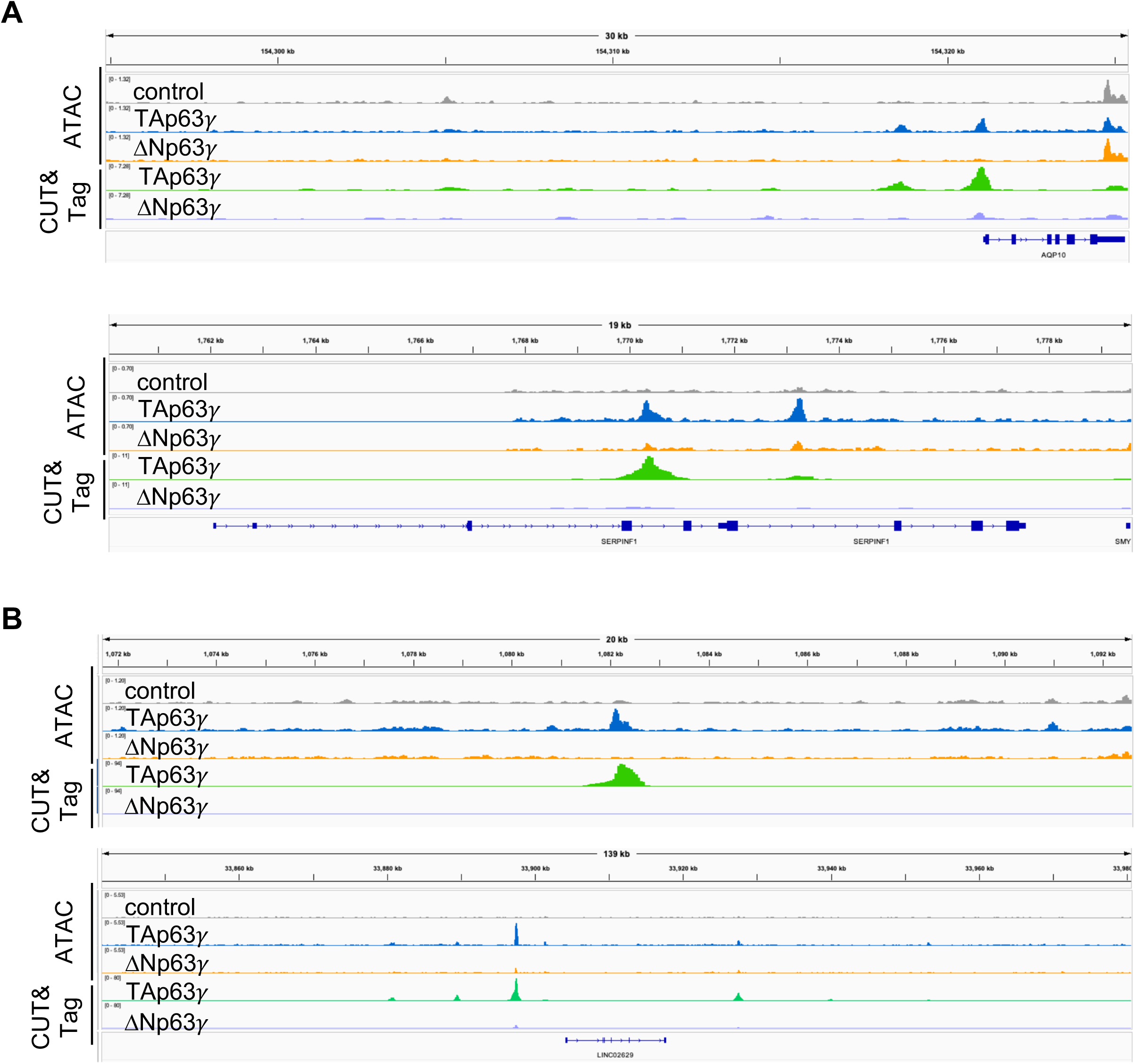
**(A/B)** Genome browser screenshots of CUT&Tag peaks and chromatin accessibility (ATAC-seq) following expression of TAp63*γ*; and ΔNp63*γ*;.

## REFERENCES

1. Lambert, S.A., Jolma, A., Campitelli, L.F., Das, P.K., Yin, Y., Albu, M., Chen, X., Taipale, J., Hughes, T.R. and Weirauch, M.T. (2018) The Human Transcription Factors. Cell, 172, 650–665.

2. Lambourne, L., Mattioli, K., Santoso, C., Sheynkman, G., Inukai, S., Kaundal, B., Berenson, A., Spirohn-Fitzgerald, K., Bhattacharjee, A., Rothman, E., et al. (2025) Widespread variation in molecular interactions and regulatory properties among transcription factor isoforms. Mol. Cell, 85, 1445–1466.e13.

3. Yang, A., Schweitzer, R., Sun, D., Kaghad, M., Walker, N., Bronson, R.T., Tabin, C., Sharpe, A., Caput, D., Crum, C., et al. (1999) p63 is essential for regenerative proliferation in limb, craniofacial and epithelial development. Nature, 398, 714–718.

4. Mills, A.A., Zheng, B., Wang, X.-J., Vogel, H., Roop, D.R. and Bradley, A. (1999) p63 is a p53 homologue required for limb and epidermal morphogenesis. Nature, 398, 708–713.

5. Yang, A., Kaghad, M., Wang, Y., Gillett, E., Fleming, M.D., Dötsch, V., Andrews, N.C., Caput, D. and McKeon, F. (1998) p63, a p53 Homolog at 3q27–29, Encodes Multiple Products with Transactivating, Death-Inducing, and Dominant-Negative Activities. Mol Cell, 2, 305–316.

6. Romano, R.-A., Smalley, K., Magraw, C., Serna, V.A., Kurita, T., Raghavan, S. and Sinha, S. (2012) ΔNp63 knockout mice reveal its indispensable role as a master regulator of epithelial development and differentiation. Development, 139, 772–782.

7. Candi, E., Rufini, A., Terrinoni, A., Dinsdale, D., Ranalli, M., Paradisi, A., Laurenzi, V.D., Spagnoli, L.G., Catani, M.V., Ramadan, S., et al. (2006) Differential roles of p63 isoforms in epidermal development: selective genetic complementation in p63 null mice. Cell Death Differ., 13, 1037–1047.

8. Su, X., Napoli, M., Abbas, H.A., Venkatanarayan, A., Bui, N.H.B., Coarfa, C., Gi, Y.J., Kittrell, F., Gunaratne, P.H., Medina, D., et al. (2017) TAp63 suppresses mammary tumorigenesis through regulation of the Hippo pathway. Oncogene, 36, 2377–2393.

9. Su, X., Chakravarti, D., Cho, M.S., Liu, L., Gi, Y.J., Lin, Y.-L., Leung, M.L., El-Naggar, A., Creighton, C.J., Suraokar, M.B., et al. (2010) TAp63 suppresses metastasis through coordinate regulation of Dicer and miRNAs. Nature, 467, 986–990.

10. Guo, X., Keyes, W.M., Papazoglu, C., Zuber, J., Li, W., Lowe, S.W., Vogel, H. and Mills, A.A. (2009) TAp63 induces senescence and suppresses tumorigenesis in vivo. Nat Cell Biol, 11, 1451–1457.

11. Somerville, T.D.D., Xu, Y., Miyabayashi, K., Tiriac, H., Cleary, C.R., Maia-Silva, D., Milazzo, J.P., Tuveson, D.A. and Vakoc, C.R. (2018) TP63-Mediated Enhancer Reprogramming Drives the Squamous Subtype of Pancreatic Ductal Adenocarcinoma. Cell Reports, 25, 1741–1755.e7.

12. Keyes, W.M., Pecoraro, M., Aranda, V., Vernersson-Lindahl, E., Li, W., Vogel, H., Guo, X., Garcia, E.L., Michurina, T.V., Enikolopov, G., et al. (2011) ΔNp63α is an oncogene that targets chromatin remodeler Lsh to drive skin stem cell proliferation and tumorigenesis. Cell Stem Cell, 8, 164–76.

13. Campbell, J.D., Yau, C., Bowlby, R., Liu, Y., Brennan, K., Fan, H., Taylor, A.M., Wang, C., Walter, V., Akbani, R., et al. (2018) Genomic, Pathway Network, and Immunologic Features Distinguishing Squamous Carcinomas. Cell Reports, 23, 194–212.e6.

14. Marshall, C.B., Beeler, J.S., Lehmann, B.D., Gonzalez-Ericsson, P., Sanchez, V., Sanders, M.E., Boyd, K.L. and Pietenpol, J.A. (2021) Tissue-specific expression of p73 and p63 isoforms in human tissues. Cell Death Dis, 12, 745.

15. McCann, A.A. and Sammons, M.A. (2025) Differential Transcriptional Activity of ΔNp63β Is Encoded by an Isoform-Specific C-Terminus. Mol. Cell. Biol., 45, 369–385.

16. Baniulyte, G., McCann, A.A., Woodstock, D.L. and Sammons, M.A. (2024) Crosstalk between paralogs and isoforms influences p63-dependent regulatory element activity. Nucleic Acids Res., 52, 13812–13831.

17. Su, X., Chakravarti, D. and Flores, E.R. (2013) p63 steps into the limelight: crucial roles in the suppression of tumorigenesis and metastasis. Nat Rev Cancer, 13, 136–143.

18. Pitzius, S., Osterburg, C., Gebel, J., Tascher, G., Schäfer, B., Zhou, H., Münch, C. and Dötsch, V. (2019) TA*p63 and GTAp63 achieve tighter transcriptional regulation in quality control by converting an inhibitory element into an additional transactivation domain. Cell Death Dis, 10, 686.

19. Deutsch, G.B., Zielonka, E.M., Coutandin, D., Weber, T.A., Schäfer, B., Hannewald, J., Luh, L.M., Durst, F.G., Ibrahim, M., Hoffmann, J., et al. (2011) DNA Damage in Oocytes Induces a Switch of the Quality Control Factor TAp63α from Dimer to Tetramer. Cell, 144, 566–576.

20. Muzellec, B., Teleńczuk, M., Cabeli, V. and Andreux, M. (2023) PyDESeq2: a python package for bulk RNA-seq differential expression analysis. Bioinformatics, 39, btad547.

21. Love, M.I., Huber, W. and Anders, S. (2014) Moderated estimation of fold change and dispersion for RNA-seq data with DESeq2. Genome Biol, 15, 550.

22. Kaya-Okur, H.S. and Henikoff, S. (2020) Bench top CUT&Tag v3. 10.17504/protocols.io.bcuhiwt6.

23. Martin, M. (2011) Cutadapt removes adapter sequences from high-throughput sequencing reads. EMBnetJ., 17, 10–12.

24. Langmead, B. and Salzberg, S.L. (2012) Fast gapped-read alignment with Bowtie 2. Nat. Methods, 9, 357–359.

25. Li, H., Handsaker, B., Wysoker, A., Fennell, T., Ruan, J., Homer, N., Marth, G., Abecasis, G., Durbin, R. and Subgroup, 1000 Genome Project Data Processing (2009) The Sequence Alignment/Map format and SAMtools. Bioinformatics, 25, 2078–2079.

26. Ramírez, F., Ryan, D.P., Grüning, B., Bhardwaj, V., Kilpert, F., Richter, A.S., Heyne, S., Dündar, F. and Manke, T. (2016) deepTools2: a next generation web server for deep-sequencing data analysis. Nucleic acids Res., 44, W160–5.

27. Zhang, Y., Liu, T., Meyer, C.A., Eeckhoute, J., Johnson, D.S., Bernstein, B.E., Nusbaum, C., Myers, R.M., Brown, M., Li, W., et al. (2008) Model-based analysis of ChIP-Seq (MACS). Genome Biol, 9, R137.

28. Quinlan, A.R. and Hall, I.M. (2010) BEDTools: a flexible suite of utilities for comparing genomic features. Bioinformatics, 26, 841–842.

29. Heinz, S., Benner, C., Spann, N., Bertolino, E., Lin, Y.C., Laslo, P., Cheng, J.X., Murre, C., Singh, H. and Glass, C.K. (2010) Simple Combinations of Lineage-Determining Transcription Factors Prime cis-Regulatory Elements Required for Macrophage and B Cell Identities. Mol. Cell, 38, 576–589.

30. Corces, M.R., Trevino, A.E., Hamilton, E.G., Greenside, P.G., Sinnott-Armstrong, N.A., Vesuna, S., Satpathy, A.T., Rubin, A.J., Montine, K.S., Wu, B., et al. (2017) An improved ATAC-seq protocol reduces background and enables interrogation of frozen tissues. Nat. Methods, 14, 959–962.

31. R, S. and G, B. (2011) DiffBind: differential binding analysis of ChIP-Seq peak data.

32. Kim, D.I., Jensen, S.C., Noble, K.A., Kc, B., Roux, K.H., Motamedchaboki, K. and Roux, K.J. (2016) An improved smaller biotin ligase for BioID proximity labeling. Mol. Biol. Cell, 27, 1188–1196.

33. Erde, J., Loo, R.R.O. and Loo, J.A. (2014) Enhanced FASP (eFASP) to Increase Proteome Coverage and Sample Recovery for Quantitative Proteomic Experiments. J. Proteome Res., 13, 1885–1895.

34. Rappsilber, J., Mann, M. and Ishihama, Y. (2007) Protocol for micro-purification, enrichment, pre-fractionation and storage of peptides for proteomics using StageTips. Nat. Protoc., 2, 1896–1906.

35. Meier, F., Brunner, A.-D., Koch, S., Koch, H., Lubeck, M., Krause, M., Goedecke, N., Decker, J., Kosinski, T., Park, M.A., et al. (2018) Online Parallel Accumulation–Serial Fragmentation (PASEF) with a Novel Trapped Ion Mobility Mass Spectrometer*. Mol. Cell. Proteom.: MCP, 17, 2534–2545.

36. Perkins, D.N., Pappin, D.J.C., Creasy, D.M. and Cottrell, J.S. (1999) Probability-based protein identification by searching sequence databases using mass spectrometry data. ELECTROPHORESIS, 20, 3551–3567.

37. Szklarczyk, D., Kirsch, R., Koutrouli, M., Nastou, K., Mehryary, F., Hachilif, R., Gable, A.L., Fang, T., Doncheva, N.T., Pyysalo, S., et al. (2023) The STRING database in 2023: protein–protein association networks and functional enrichment analyses for any sequenced genome of interest. Nucleic Acids Res., 51, D638–D646.

38. Lotthammer, J.M., Hernández-García, J., Griffith, D., Weijers, D., Holehouse, A.S. and Emenecker, R.J. (2024) Metapredict enables accurate disorder prediction across the Tree of Life. bioRxiv, 10.1101/2024.11.05.622168.

39. Enthart, A., Klein, C., Dehner, A., Coles, M., Gemmecker, G., Kessler, H. and Hagn, F. (2016) Solution structure and binding specificity of the p63 DNA binding domain. Sci Rep-uk, 6, 26707.

40. Jumper, J., Evans, R., Pritzel, A., Green, T., Figurnov, M., Ronneberger, O., Tunyasuvunakool, K., Bates, R., Žídek, A., Potapenko, A., et al. (2021) Highly accurate protein structure prediction with AlphaFold. Nature, 596, 583–589.

41. Fleming, J., Magana, P., Nair, S., Tsenkov, M., Bertoni, D., Pidruchna, I., Afonso, M.Q.L., Midlik, A., Paramval, U., Žídek, A., et al. (2025) AlphaFold Protein Structure Database and 3D-Beacons: New Data and Capabilities. J. Mol. Biol., 437, 168967.

42. Ginell, G.M., Emenecker, R.J., Lotthammer, J.M., Keeley, A.T., Plassmeyer, S.P., Razo, N., Usher, E.T., Pelham, J.F. and Holehouse, A.S. (2025) Sequence-based prediction of intermolecular interactions driven by disordered regions. Science, 388, eadq8381.

43. Joseph, J.A., Reinhardt, A., Aguirre, A., Chew, P.Y., Russell, K.O., Espinosa, J.R., Garaizar, A. and Collepardo-Guevara, R. (2021) Physics-driven coarse-grained model for biomolecular phase separation with near-quantitative accuracy. Nat. Comput. Sci., 1, 732–743.

44. Meer, D. van der, Barthorpe, S., Yang, W., Lightfoot, H., Hall, C., Gilbert, J., Francies, H.E. and Garnett, M.J. (2019) Cell Model Passports—a hub for clinical, genetic and functional datasets of preclinical cancer models. Nucleic Acids Res., 47, D923–D929.

45. Heo, D.S., Snyderman, C., Gollin, S.M., Pan, S., Walker, E., Deka, R., Barnes, E.L., Johnson, J.T., Herberman, R.B. and Whiteside, T.L. (1989) Biology, cytogenetics, and sensitivity to immunological effector cells of new head and neck squamous cell carcinoma lines. Cancer Res., 49, 5167–75.

46. Krauskopf, K., Gebel, J., Kazemi, S., Tuppi, M., Löhr, F., Schäfer, B., Koch, J., Güntert, P., Dötsch, V. and Kehrloesser, S. (2018) Regulation of the Activity in the p53 Family Depends on the Organization of the Transactivation Domain. Structure, 26, 1091–1100.e4.

47. Coutandin, D., Osterburg, C., Srivastav, R.K., Sumyk, M., Kehrloesser, S., Gebel, J., Tuppi, M., Hannewald, J., Schäfer, B., Salah, E., et al. (2016) Quality control in oocytes by p63 is based on a spring-loaded activation mechanism on the molecular and cellular level. Elife, 5, e13909.

48. Strasser, A., Harris, A.W., Jacks, T. and Cory, S. (1994) DNA damage can induce apoptosis in proliferating lymphoid cells via p53-independent mechanisms inhibitable by Bcl-2. Cell, 79, 329–339.

49. Chiou, S.K., Rao, L. and White, E. (1994) Bcl-2 blocks p53-dependent apoptosis. Mol. Cell. Biol., 14, 2556–2563.

50. Helton, E.S., Zhu, J. and Chen, X. (2006) The Unique NH2-terminally Deleted (ΔN) Residues, the PXXP Motif, and the PPXY Motif Are Required for the Transcriptional Activity of the ΔN Variant of p63*. J Biol Chem, 281, 2533–2542.

51. Ptashne, M. (1988) How eukaryotic transcriptional activators work. Nature, 335, 683–689.

52. Staller, M.V., Ramirez, E., Kotha, S.R., Holehouse, A.S., Pappu, R.V. and Cohen, B.A. (2022) Directed mutational scanning reveals a balance between acidic and hydrophobic residues in strong human activation domains. Cell Syst, 13, 334–345.e5.

53. DelRosso, N., Tycko, J., Suzuki, P., Andrews, C., Aradhana, Mukund, A., Liongson, I., Ludwig, C., Spees, K., Fordyce, P., et al. (2023) Large-scale mapping and mutagenesis of human transcriptional effector domains. Nature, 616, 365–372.

54. Kotha, S.R. and Staller, M.V. (2023) Clusters of acidic and hydrophobic residues can predict acidic transcriptional activation domains from protein sequence. GENETICS, 225, iyad131.

55. Sigler, P.B. (1988) Transcriptional activation. Acid blobs and negative noodles. Nature, 333, 210–2.

56. Ferrie, J.J., Karr, J.P., Tjian, R. and Darzacq, X. (2022) “Structure”-function relationships in eukaryotic transcription factors: The role of intrinsically disordered regions in gene regulation. Mol Cell, 82, 3970–3984.

57. Holehouse, A.S. and Kragelund, B.B. (2023) The molecular basis for cellular function of intrinsically disordered protein regions. Nat. Rev. Mol. Cell Biol., 10.1038/s41580-023-00673-0.

58. Jonas, F., Navon, Y. and Barkai, N. (2025) Intrinsically disordered regions as facilitators of the transcription factor target search. Nat. Rev. Genet., 10.1038/s41576-025-00816-3.

59. Chen, Y., Cattoglio, C., Dailey, G.M., Zhu, Q., Tjian, R. and Darzacq, X. (2022) Mechanisms governing target search and binding dynamics of hypoxia-inducible factors. eLife, 11, e75064.

60. Brodsky, S., Jana, T., Mittelman, K., Chapal, M., Kumar, D.K., Carmi, M. and Barkai, N. (2020) Intrinsically Disordered Regions Direct Transcription Factor In Vivo Binding Specificity. Mol Cell, 79, 459–471.e4.

61. Aubrey, B.J., Kelly, G.L., Janic, A., Herold, M.J. and Strasser, A. (2018) How does p53 induce apoptosis and how does this relate to p53-mediated tumour suppression? Cell Death Differ, 25, 104–113.

62. Kerr, J.B., Hutt, K.J., Michalak, E.M., Cook, M., Vandenberg, C.J., Liew, S.H., Bouillet, P., Mills, A., Scott, C.L., Findlay, J.K., et al. (2012) DNA Damage-Induced Primordial Follicle Oocyte Apoptosis and Loss of Fertility Require TAp63-Mediated Induction of Puma and Noxa. Mol. Cell, 48, 343–352.

63. Bolcun-Filas, E., Rinaldi, V.D., White, M.E. and Schimenti, J.C. (2014) Reversal of Female Infertility by Chk2 Ablation Reveals the Oocyte DNA Damage Checkpoint Pathway. Science, 343, 533–536.

64. Lin-Shiao, E., Lan, Y., Coradin, M., Anderson, A., Donahue, G., Simpson, C.L., Sen, P., Saffie, R., Busino, L., Garcia, B.A., et al. (2018) KMT2D regulates p63 target enhancers to coordinate epithelial homeostasis. Gene Dev, 32, 181–193.

65. LeBoeuf, M., Terrell, A., Trivedi, S., Sinha, S., Epstein, J.A., Olson, E.N., Morrisey, E.E. and Millar, S.E. (2010) Hdac1 and Hdac2 Act Redundantly to Control p63 and p53 Functions in Epidermal Progenitor Cells. Dev. Cell, 19, 807–818.

66. Ramsey, M.R., He, L., Forster, N., Ory, B. and Ellisen, L.W. (2011) Physical Association of HDAC1 and HDAC2 with p63 Mediates Transcriptional Repression and Tumor Maintenance in Squamous Cell Carcinoma. Cancer Res., 71, 4373–4379.

67. Grant, P.A., Duggan, L., Côté, J., Roberts, S.M., Brownell, J.E., Candau, R., Ohba, R., Owen-Hughes, T., Allis, C.D., Winston, F., et al. (1997) Yeast Gcn5 functions in two multisubunit complexes to acetylate nucleosomal histones: characterization of an Ada complex and the SAGA (Spt/Ada) complex. Genes Dev., 11, 1640–1650.

68. Herbst, D.A., Esbin, M.N., Louder, R.K., Dugast-Darzacq, C., Dailey, G.M., Fang, Q., Darzacq, X., Tjian, R. and Nogales, E. (2021) Structure of the human SAGA coactivator complex. Nat. Struct. Mol. Biol., 28, 989–996.

69. Timmers, H.Th.M. (2021) SAGA and TFIID: Friends of TBP drifting apart. Biochim. Biophys. Acta (BBA) - Gene Regul. Mech., 1864, 194604.

70. Bassi, Z.I., Fillmore, M.C., Miah, A.H., Chapman, T.D., Maller, C., Roberts, E.J., Davis, L.C., Lewis, D.E., Galwey, N.W., Waddington, K.E., et al. (2018) Modulating PCAF/GCN5 Immune Cell Function through a PROTAC Approach. ACS Chem. Biol., 13, 2862–2867.

71. Nagy, Z. and Tora, L. (2007) Distinct GCN5/PCAF-containing complexes function as co-activators and are involved in transcription factor and global histone acetylation. Oncogene, 26, 5341–5357.

72. Yamauchi, T., Yamauchi, J., Kuwata, T., Tamura, T., Yamashita, T., Bae, N., Westphal, H., Ozato, K. and Nakatani, Y. (2000) Distinct but overlapping roles of histone acetylase PCAF and of the closely related PCAF-B/GCN5 in mouse embryogenesis. Proc. Natl. Acad. Sci., 97, 11303–11306.

73. Malone, C.F., Mabe, N.W., Forman, A.B., Alexe, G., Engel, K.L., Chen, Y.-J.C., Soeung, M., Salhotra, S., Basanthakumar, A., Liu, B., et al. (2024) The KAT module of the SAGA complex maintains the oncogenic gene expression program in MYCN-amplified neuroblastoma. Sci. Adv., 10, eadm9449.

74. Maia-Silva, D., Cunniff, P.J., Schier, A.C., Skopelitis, D., Trousdell, M.C., Moresco, P., Gao, Y., Kechejian, V., He, X.-Y., Sahin, Y., et al. (2024) Interaction between MED12 and ΔNp63 activates basal identity in pancreatic ductal adenocarcinoma. Nat. Genet., 10.1038/s41588-024-01790-y.

75. Mindel, V., Brodsky, S., Yung, H., Manadre, W. and Barkai, N. (2024) Revisiting the model for coactivator recruitment: Med15 can select its target sites independent of promoter-bound transcription factors. Nucleic Acids Res., 52, 12093–12111.

76. Staller, M.V. (2022) Transcription factors perform a 2-step search of the nucleus. Genetics, 10.1093/genetics/iyac111.

77. McMahon, S.B., Buskirk, H.A.V., Dugan, K.A., Copeland, T.D. and Cole, M.D. (1998) The Novel ATM-Related Protein TRRAP Is an Essential Cofactor for the c-Myc and E2F Oncoproteins. Cell, 94, 363–374.

78. Chesnutt, K.V., Yayli, G., Toelzer, C., Damilot, M., Cox, K., Gautam, G., Berger, I., Tora, L. and Poirier, M.G. (2024) ATAC and SAGA histone acetyltransferase modules facilitate transcription factor binding to nucleosomes independent of their acetylation activity. Nucleic Acids Res., 53, gkae1120.

79. Wang, H., Dienemann, C., Stützer, A., Urlaub, H., Cheung, A.C.M. and Cramer, P. (2020) Structure of the transcription coactivator SAGA. Nature, 577, 717–720.

80. Baptista, T., Grünberg, S., Minoungou, N., Koster, M.J.E., Timmers, H.T.M., Hahn, S., Devys, D. and Tora, L. (2017) SAGA Is a General Cofactor for RNA Polymerase II Transcription. Mol. Cell, 68, 130–143.e5.

81. Papai, G., Frechard, A., Kolesnikova, O., Crucifix, C., Schultz, P. and Ben-Shem, A. (2020) Structure of SAGA and mechanism of TBP deposition on gene promoters. Nature, 577, 711–716.

82. Neumayr, C., Haberle, V., Serebreni, L., Karner, K., Hendy, O., Boija, A., Henninger, J.E., Li, C.H., Stejskal, K., Lin, G., et al. (2022) Differential cofactor dependencies define distinct types of human enhancers. Nature, 606, 406–413.

83. Bell, C.C., Balic, J.J., Talarmain, L., Gillespie, A., Scolamiero, L., Lam, E.Y.N., Ang, C.-S., Faulkner, G.J., Gilan, O. and Dawson, M.A. (2024) Comparative cofactor screens show the influence of transactivation domains and core promoters on the mechanisms of transcription. Nat. Genet., 10.1038/s41588-024-01749-z.

84. Yang, L., Shah, M., Chavalit, T., Spek, A.M., Lim, W.F., Zhong, L., Safari, V.N., Haddad, M., Vu, T.H., Mackay, J.P., et al. (2025) WDR5 serves in co-activation and influences genome targeting of KLF3. Nucleic Acids Res., 53, gkaf977.

85. Bonnet, J., Wang, C.-Y., Baptista, T., Vincent, S.D., Hsiao, W.-C., Stierle, M., Kao, C.-F., Tora, L. and Devys, D. (2014) The SAGA coactivator complex acts on the whole transcribed genome and is required for RNA polymerase II transcription. Genes Dev., 28, 1999–2012.

86. Krebs, A.R., Karmodiya, K., Lindahl-Allen, M., Struhl, K. and Tora, L. (2011) SAGA and ATAC Histone Acetyl Transferase Complexes Regulate Distinct Sets of Genes and ATAC Defines a Class of p300-Independent Enhancers. Mol. Cell, 44, 410–423.

87. Huisinga, K.L. and Pugh, B.F. (2004) A Genome-Wide Housekeeping Role for TFIID and a Highly Regulated Stress-Related Role for SAGA in Saccharomyces cerevisiae. Mol. Cell, 13, 573–585.

88. Kremer, S.B. and Gross, D.S. (2009) SAGA and Rpd3 Chromatin Modification Complexes Dynamically Regulate Heat Shock Gene Structure and Expression*. J. Biol. Chem., 284, 32914–32931.

89. Basehoar, A.D., Zanton, S.J. and Pugh, B.F. (2004) Identification and Distinct Regulation of Yeast TATA Box-Containing Genes. Cell, 116, 699–709.

90. Suh, E.-K., Yang, A., Kettenbach, A., Bamberger, C., Michaelis, A.H., Zhu, Z., Elvin, J.A., Bronson, R.T., Crum, C.P. and McKeon, F. (2006) p63 protects the female germ line during meiotic arrest. Nature, 444, 624–628.

91. Koster, M.I., Kim, S., Mills, A.A., DeMayo, F.J. and Roop, D.R. (2004) p63 is the molecular switch for initiation of an epithelial stratification program. Gene Dev, 18, 126–131.

92. Koster, M.I., Dai, D., Marinari, B., Sano, Y., Costanzo, A., Karin, M. and Roop, D.R. (2007) p63 induces key target genes required for epidermal morphogenesis. Proc National Acad Sci, 104, 3255–3260.

